# Exogenous Na_V_1.1 activity in excitatory and inhibitory neurons reverts Dravet syndrome comorbidities when delivered post-symptom onset in mice with Dravet

**DOI:** 10.1101/2022.06.10.495591

**Authors:** Saja Fadila, Bertrand Beucher, Iria González-Dopeso Reyes, Anat Mavashov, Marina Brusel, Karen Anderson, Ethan M Goldberg, Ana Ricobaraza, Ruben Hernandez-Alcoceba, Eric J Kremer, Moran Rubinstein

**Affiliations:** Goldschleger Eye Research Institute, Faculty of Medicine, Tel Aviv University, Tel Aviv, Israel; Department of Human Molecular Genetics and Biochemistry, Faculty of Medicine, Tel Aviv University, Tel Aviv, Israel; PVM, BioCampus, CNRS, INSERM, University of Montpellier, Montpellier, France; Institut de Génétique Moléculaire de Montpellier, Université de Montpellier, CNRS, Montpellier, France; Sagol School of Neuroscience, Tel Aviv University, Tel Aviv, Israel; Department of Neuroscience, The Perelman School of Medicine at The University of Pennsylvania, Philadelphia, USA; Gene Therapy and Regulation of Gene Expression Program, CIMA, University of Navarra. IdiSNA, Navarra Institute for Health Research, Pamplona, Spain

## Abstract

Dravet syndrome (DS), an intractable childhood epileptic encephalopathy with a high fatality rate, is caused by loss-of-function mutations in one allele of *SCN1A*, which encodes Na_V_1.1. In contrast to other epilepsies, pharmaceutical treatment for DS is limited. Here, we demonstrate that viral vector-mediated delivery of a codon-modified *SCN1A* cDNA improves DS comorbidities in juvenile and adolescent DS mice (*Scn1a*^A1783V/WT^). Notably, bilateral vector injections into the hippocampus or thalamus of DS mice improved the survival of the mice, reduced the occurrence of epileptic spikes, provided protection from thermally-induced seizures, and corrected background electrocorticography activity. Together, our results provide a proof-of-concept for the potential of *SCN1A* delivery as a therapeutic approach for infants and adolescents with DS-associated comorbidities.

## Introduction

Dravet syndrome (DS) is a rare and severe form of developmental epileptic encephalopathy (DEE). Infants with DS appear to develop normally during the first six months of life, but subsequently start to exhibit febrile seizures. During the following months, recurrent refractory spontaneous seizures become increasingly more frequent and global developmental delays begin. During the early school years, which represent the chronic phase of the disease, the frequency of seizures declines, but the non-epileptic comorbidities persist (1–3).

The vast majority of DS cases are caused by *de novo* mutations in one copy of *SCN1A*. These cause loss-of-function (LoF) and therefore result in haploinsufficiency in the activity of the voltage-gated sodium channel, Na_V_1.1 (4). Thus, a strategy in which Na_V_1.1 channels are reintroduced into the central nervous system (CNS) neurons, can represent a useful therapeutic approach, regardless of the underlying *SCN1A* mutation.

One option for gene therapy is to provide exogenous Na_V_1.1 via delivery of the *SCN1A* cDNA. However, due to the size of the coding region (∼6 kbp), commonly used vectors, such as adeno-associated virus (AAV), are poorly adapted. In an attempt to overcome the “size” obstacle, previous gene therapy strategies for DS have involved (*i*) enhanced expression of the endogenous *Scn1a* via transcriptional activation (5–7); (*ii*) overexpression of *SCN1B,* which encodes Na_V_β1, an auxiliary subunit that increase Na_V_1.1 channel complex efficacy (8); and, (*iii*) antisense oligonucleotide-mediated downregulation of *SCN8A* (9). While these approaches showed proof of principle in mouse models of DS, the therapeutic potential was manifested only when administered soon after birth, during the asymptomatic, pre-epileptic stage (5, 7–9). As clinical diagnoses are rarely confirmed prior to progression to recurrent spontaneous seizures, therapies able to improve the epilepsy and protect from sudden unexpected death in epilepsy (SUDEP), following the onset of severe intractable seizures, remain a critical unmet need. Recent studies indicate that activation of the *Scn1a* gene in adult mice can reverse DS (10). While this required reactivation of a floxed stopped *Scn1a* with Cre recombinase, and therefore is not directly translatable for clinical use, these results provide a validation that restoration of Na_V_1.1 activity, in critical brain regions, can ameliorate DS symptoms.

In contrast to AAV, helper-dependent (HD) adenovirus vectors can harbour up to 37 kb of exogenous sequence and allow transgene expression almost immediately post-injection (11–13). The large cloning capacity allows the delivery of multiple cassettes with large transcription regulatory sequences. Moreover, canine adenovirus type 2 (CAV-2) vectors offer substantial advantages for gene transfer to the CNS (13–15). First, by preferentially using the coxsackievirus and adenovirus receptor (CAR) as an attachment molecule, whose expression is found primarily by neurons in the brain parenchyma (16), CAV-2 vectors preferentially transduce neurons (17). Second, with its robust retrograde axonal transport, local injections lead to expression across connected brain regions (17). Third, like AAV vectors, injection of both E1/E3-deleted and HD CAV-2 vectors generate long-term transgene expression in the CNS of rodents (15, 18).

Here, we demonstrate that CAV-2-mediated CNS delivery of a codon-modified *SCN1A* cDNA in mice with DS significantly improves the survival of juvenile mice, reduces the occurrence of spontaneous seizures and epileptic spike frequency, and increases the temperature threshold for febrile seizures in juvenile and adolescent mice. Together, our results demonstrate that neuronal delivery of an *SCN1A* expression cassette is a promising therapeutic approach for DS.

## Results

### Transcriptionally-targeted transgene expression

The current dogma is that many DS-associated symptoms are caused by inhibitory neurons dysfunction (19). However, others have postulated that reduced Na_V_1.1 activity in excitatory neurons may also be associated with DS comorbidities (20–22). Moreover, hippocampal dysfunction is thought to play a key role in DS pathophysiology (23–25). We therefore initiated assays to determine whether we could selectively target expression to different neuronal populations in the hippocampus. Because direct selective infection of these neuronal subtypes is not yet technically feasible, we opted for transcriptional control of the expression cassette. To this end, we prepared CAV-2 vectors containing a fluorescent protein driven by (*i*) a strong non-specific CAG promoter (cytomegalovirus enhancer, chicken β-actin promoter and rabbit β-globin splice acceptor site) that should drive expression in all vector-transduced cells; (*ii*) a human synapsin promoter (hSyn), which drives transgene expression in most neuronal populations; (*iii*) a neuron-specific enolase (NSE) promoter, which drives moderate levels of transgene expression in excitatory and inhibitory neurons (26) and (*iv*) a Dlx5/6 promoter (Dlx5 and Dlx6 encode two homeobox transcription factors expressed by developing and mature GABAergic interneurons), which, when incorporated into a viral vector, can lead to preferential expression in inhibitory neurons (27, 28).

When these vectors were injected into the hippocampus of adult mice the pattern of expression, as expected, depended on the promoter. The CAG promoter generated robust and widespread transgene expression at the site of injection and, due to the retrograde transport of CAV-2, was also expressed in excitatory neurons from the subiculum and multiple neocortical layers that project into the hippocampus (**Fig. 1A-D**). Although at lower levels, the hSyn promoter also led to widespread transgene expression at the site of injection and in afferent regions (**Fig. 1E-H**). The NSE promoter (**Fig. 1I-L**) resulted in an intermediate level of expression in excitatory and inhibitory neurons around the injection sites (based on location & morphology), as well as in additional hippocampal projecting regions in the cortex and thalamus (**Fig. 1I**). Conversely, the Dlx5/6 promoter produced a localized transgene expression principally in hippocampal interneurons (based on location & morphology) (**Fig. 1M-P**). After consideration of the expression patterns in excitatory and inhibitory neurons, biodistribution, and the probable requirement for moderate levels of Na_V_1.1, we opted for the pan-neuronal expression mediated by the NSE promoter to drive SCN1A expression by CAV-2 vectors.

**Fig. 1.**
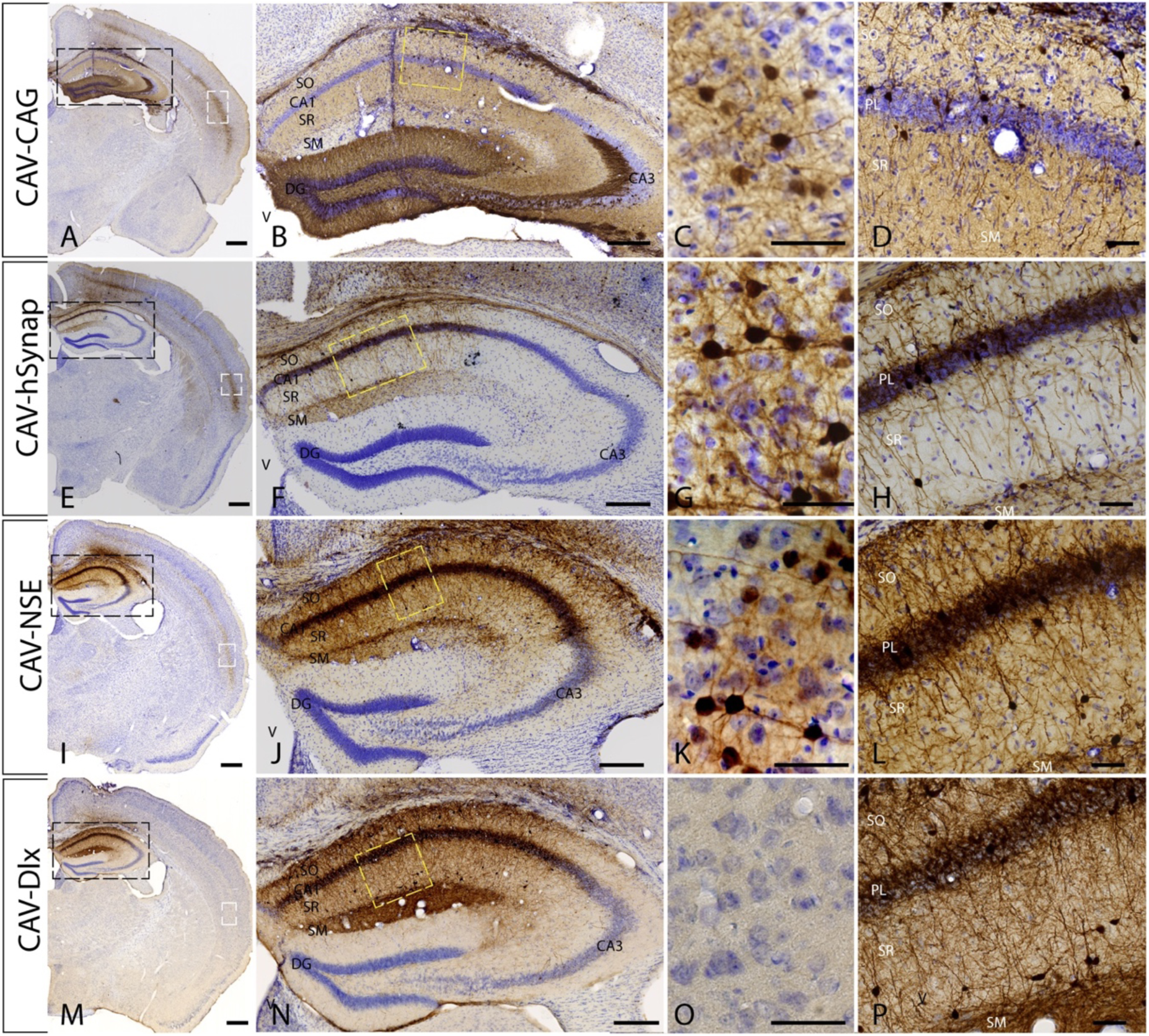
Transcriptional control of transgene expression in CAV-2 vectors following injection into the mouse hippocampus. CAV-2 vectors containing variable promoters upstream of an mCitrine open reading frame were generated. 1 x 10^9^ physical particles of each vector were injected bilaterally into the hippocampus of adult mice. Background staining is cresyl violet. mCitrine expression is shown by immunohistology DAB staining (dark brown). (**A-D**) CAG promoter driven mCitrine expression: in this mouse, the vector was deposited in the DG. Expression can be seen further up the needle track and into the CA1 region (**E-H**) hSyn driven mCitrine expression: in this mouse, the vector was deposited in the CA1 region (**I-L**) NSE driven mCitrine expression: in this mouse, the vector was deposited in the CA1 region; and (**M-P**) Dlx driven mCitrine expression: in this mouse, the vector was deposited in the CA1 region. The left-hand column shows one hemisphere, the second column shows the hippocampus (magnification of black box in the first column), the third column shows transgene expression in the neocortex (magnification of white box in the first column), and the fourth column shows transgene expression in the CA1 (magnification of yellow box in the second column). Scale bar: A,E,I and M: 1 mm; B,F,J,N: 250 μm, C,G,K and O: 10 μm, D,H,L and P: 50 μm.

### CAV-SCN1A-mediated Na_V_1.1 activity

In addition to the size of the *SCN1A* cDNA, the sequence is prone to rearrangement when subcloned into a plasmid and propagated in *E. coli* (19). However, codon modification can enhance the stability and thereby facilitate the creation of vectors (29). To be able to readily subclone the *SCN1A* cDNA, we generated a codon-modified open reading frame and then manually screened for and eliminated short (10-15 bp) repeat sequences found primarily in the regions encoding the transmembrane domains of Na_V_1.1. Because detection of exogenous Na_V_1.1 by immunohistology could be problematic, a C-terminal hemagglutinin (HA) tag was incorporated into Na_V_1.1 to allow detection of vector-mediated expression (versus endogenous protein).

CAV-2 vectors containing the NSE promoter, driving expression of the codon-modified *SCN1A* cDNA (CAV-SCN1A), or a control vector (CAV-GFP) were generated and purified as previously described (30) (**Fig. S1**). To confirm the generation of functional Na_V_1.1 channels, we incubated CAV-SCN1A with DK cells (30) and recorded robust voltage-gated sodium currents with the biophysical properties that are characteristic of Na_V_1.1 (**Fig. 2A, B**) (31). Of note, comparing vectors with and without the HA tag demonstrated that this tag does not affect the biophysical properties of the channel (**Fig. 2C, D**).

**Fig. 2.**
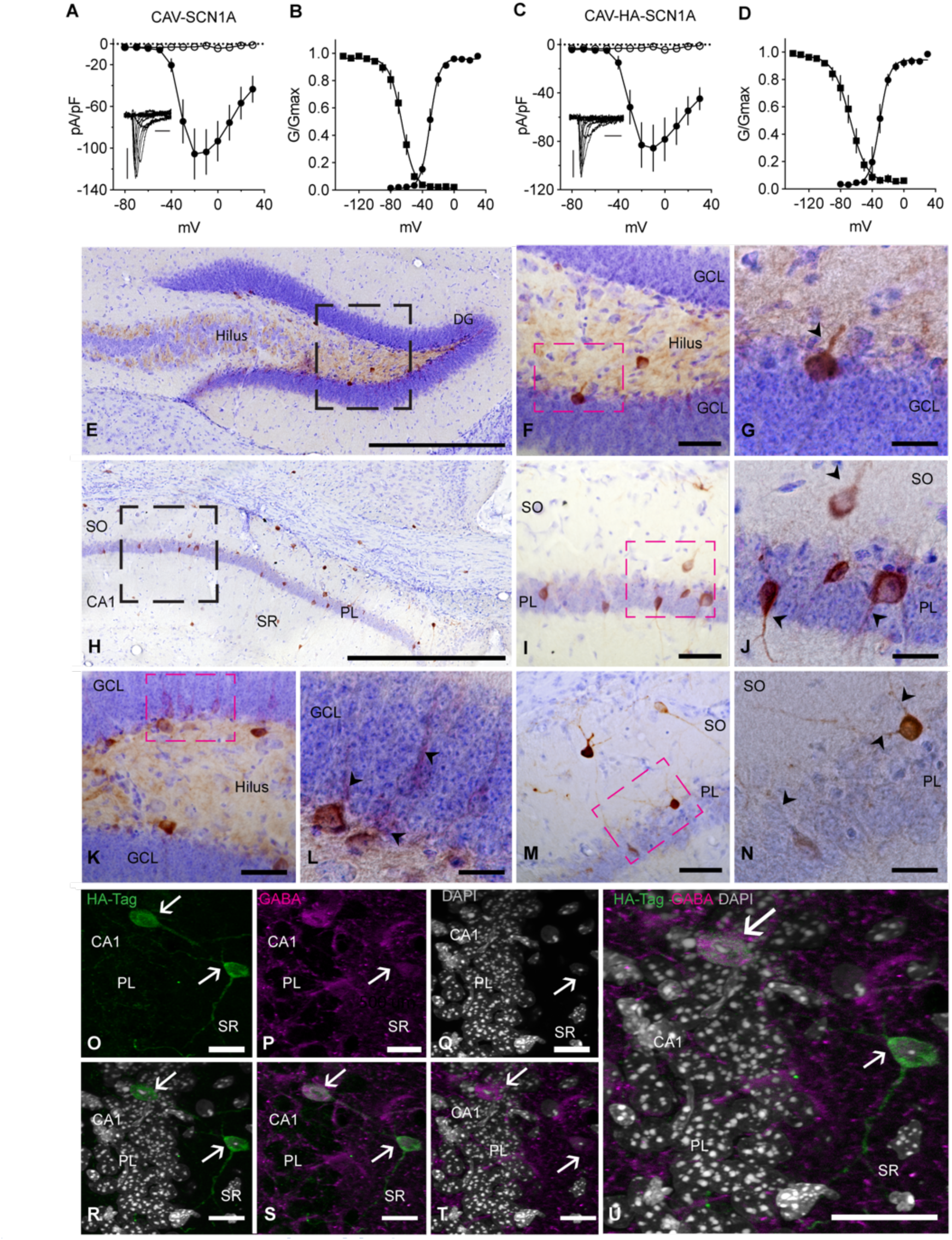
Functional expression of the SCN1A transgene. (**A-B**) Voltage current relationship (A) and the biophysical properties (B) of sodium currents following expression of CAV-SCN1A in DK cells. The half voltage of activation was -30.4 ± 0.7 mV; the half voltage of inactivation was -65.73 ± 0.9 mV. (**C-D**) Voltage current relationship and the biophysical properties of sodium currents following expression of CAV-HA-SCN1A in DK cells. The half voltage of activation was -30.6 ± 1.2 mV and half voltage of inactivation was -67.8 ± 1.8 mV. Inset in A and C: a representative sodium current traces, calibrators: 500 pA, 2 ms. The empty symbols depict the current recorded from DK cells infected with the control CAV-GFP control vector (n = 10), closed symbols depict CAV-SCN1A (n = 9) or CAV-HA-SCN1A (n = 7), as indicated. (**E-U**) CAV-HA-SCN1A. 1 x 10^9^ physical particles were injected bilaterally into the hippocampus of adult mice. (**E-N**) Background staining is cresyl violet. HA expression is shown by immunohistology DAB staining (dark brown). Approximately 1,000 HA-immunoreactive cells/mouse, with neuronal morphology could be readily identified. (**E, H**) HA immunoreactivity in the hippocampus, scale bar 500 μm. (**F, I**) magnification of the black box in E and H, and additional examples (**K, M**) scale bar 50 μm. (**G, J, L, N**) Magnification of the red box, scale bar 20 μm. (**O-U**) Immunofluorescence in a mouse injected with CAV-HA-SCN1A. (**O**) HA immunoreactivity (IR), (**P**) GABA IR, (**Q**) DAPI IR and merge (**R-U**). Scale bars 20 µm.

Following injection of CAV-SCN1A into the hippocampus of adult mice, HA-immunoreactivity was detected in the projections and soma of hippocampal neurons, some of which were GABA immunoreactive (**Fig. 2E-U**). Of note, the differences in the detection and overall distribution of HA-SCN1A (**Fig. 2E-P**), compared to that of GFP (**Fig. 1I-L**), are probably an underestimation of HA-SCN1A expression, which is due to the location of HA-Na_V_1.1 in the membrane (vs. cytoplasm for GFP), the difficulties in detection of Na_V_ channels in fixed tissue (32, 33), and the use of a monoclonal anti-HA vs. polyclonal anti-GFP antibodies.

### CAV-SCN1A injections revert epileptic phenotypes in adolescent DS mice

There are multiple mouse models of DS that faithfully reproduce the hallmarks of DS pathology (20, 34–41). Most “DS mice” display age-dependent progression of the severity of the epilepsy with spontaneous seizures that begin around postnatal day (P) 18, and premature death that peaks during the 4^th^ week of life. Surviving DS mice enter a chronic stage, in which the frequency of spontaneous convulsive seizures, and mortality are reduced (38). The genetic background dramatically affects the severity of the epileptic phenotype in DS mice (21, 35, 42, 43). Here, we used one of the more demanding DS mouse models, harboring a missense mutation (*Scn1a*^A1783V/WT^) on the C57BL/6J background. The *Scn1a*^A1783V^ mutation causes LoF of Na_V_1.1 (44), recapitulating the characteristic neuronal alterations of DS (44–46). All DS *Scn1a*^A1783V/WT^ mice experience spontaneous seizures, with an overall mortality rate of over 50% (38, 41, 45). However, unlike other models with *Scn1a* truncation mutations, the *Scn1a*^A1783V^ does not affect the overall *Scn1a* mRNA or total Na_V_1.1 levels (41).

Recently, vector-mediated expression of Na_V_1.1, was shown to reduce DS pathology in adolescent DS *Scn1a*^A1783V/WT^ mice. Mora-Jimenez et al. (2021) combined bilateral injections into six locations throughout the cortex, basal ganglia, and cerebellum. However, Na_V_1.1 immunoreactivity was limited to the area flanking the needle track. Considering the robust retrograde transport of CAV-2 vectors, we postulated that injections into the hippocampi may be sufficient to improve the epilepsy in adolescent DS mice. To that end, bilateral injections of CAV-SCN1A or CAV-GFP were performed in 5-week-old healthy (WT) and DS *Scn1a*^A1783V/WT^ mice. As already described, the prevalence of SUDEP is low after P30, during the chronic stage of DS (35, 36, 38, 41). Inasmuch, only one mock-treated DS mouse died out of the cohorts of mock (CAV-GFP injected) and CAV-SCN1A treated mice (**Fig. S2**). In lieu of survival, we used electrocorticography (ECoG) to examine the impact of CAV-SCN1A on aberrant neuronal activity. Two weeks post-injection, we implanted depth electrodes in the hippocampus, and intracranial electrodes in the somatosensory cortex for ECoG recordings. Initially, these recordings showed that CAV-SCN1A injections in WT mice did not induce aberrant neuronal activity (**Fig. 3A, C**). Moreover, in DS mice, hippocampi CAV-SCN1A injections reduced the occurrence of epileptic spikes (**Fig. 3A-D**).

**Fig. 3.**
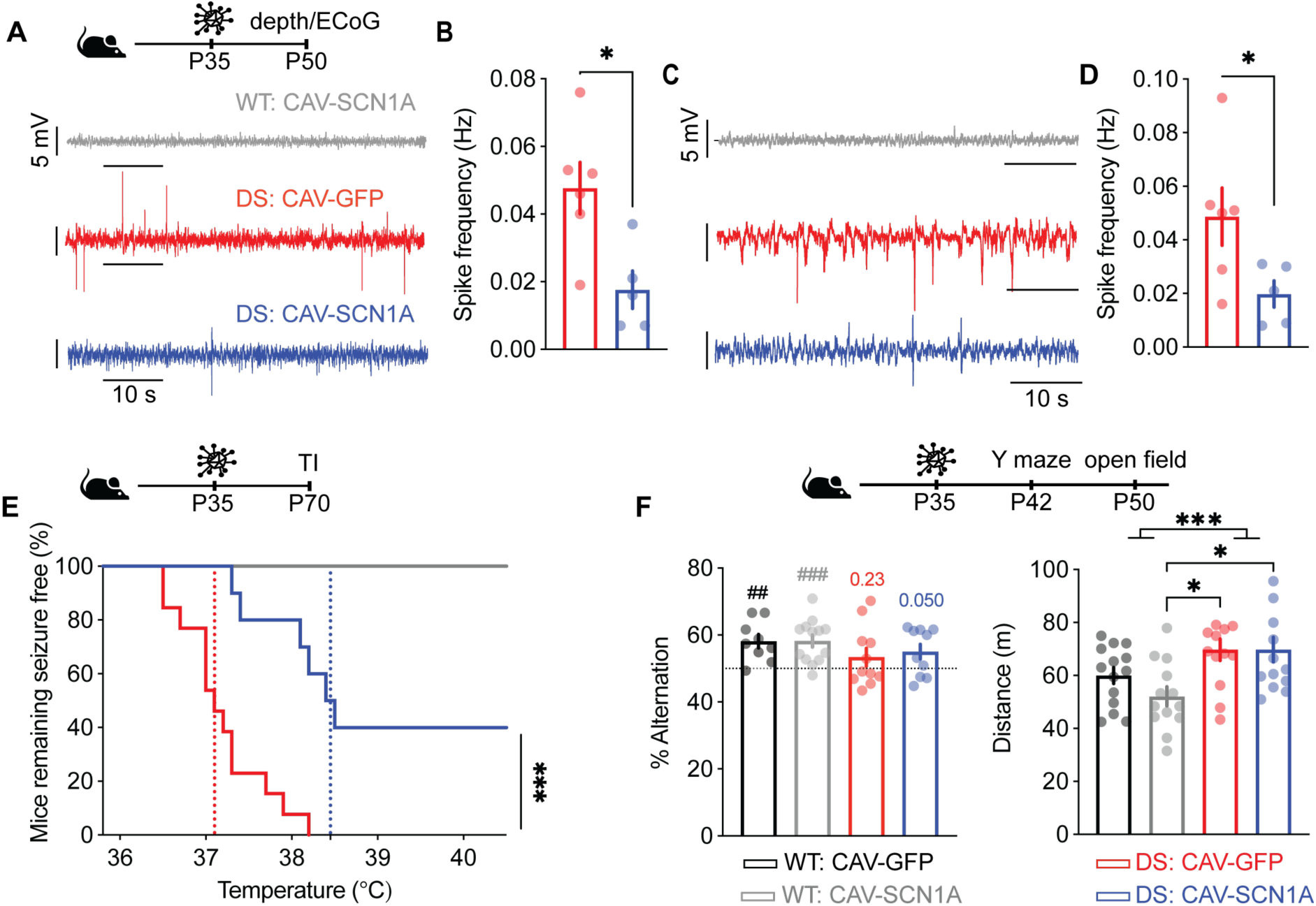
Hippocampal injections of CAV-SCN1A during the chronic stage of DS reduce the epilepsy symptoms. CAV-GFP or CAV-SCN1A (1 × 10^9^ physical particles) were injected bilaterally into the hippocampus of 5-week-old WT and DS mice. (**A-B**) Two weeks after the treatments, depth electrodes were implanted into the hippocampus, at the site of injection. Example traces (**A**) and quantification of the spike frequencies (**B**). Epileptic activity was not detected in WT mice injected with CAV-SCN1A (n = 3); DS: CAV-GFP (n = 6); DS: CAV-SCN1A (n = 5). **(C-D)** Example traces of cortical ECoG recordings (**C**) and quantification of the spike frequencies (**D**). WT: CAV-SCN1A (n = 2); DS: CAV-GFP (n = 6); DS: CAV-SCN1A (n = 5). (**E**) Mice remaining free of thermally-induced (TI) seizures. The dotted lines represent the median seizure temperature. WT: CAV-GFP, n = 11; WT: CAV-SCN1A, n = 7; DS: CAV-GFP, n = 13; DS: CAV-SCN1A, n = 10. See **Fig. S3** for separate analyses of males and females. (**F**) Left: Spontaneous alternations in the Y maze. The dotted line signifies the chance level, expected from random alternation. The markings above the bars indicate statistical analysis using one-sample *t* test relative to 50%. WT: CAV-GFP (n = 9); WT: CAV-SCN1A (n = 13); DS: CAV-GFP (n = 11); DS: CAV-SCN1A (n = 10). Right: The distance moved in the open field. WT: CAV-GFP (n = 14); WT: CAV-SCN1A (n = 13); DS: CAV-GFP (n = 13); DS: CAV-SCN1A (n = 13). See **Fig. S4**, for gender separated analyses. See **Table S1** for details about statistical analyses.

Another hallmark of DS is the sensitivity to thermally-induced seizures. Anti-seizure medications, which are effective in treating patients with DS, can elevate the threshold of thermally-induced seizures in DS mice (47, 48). Strikingly, ∼40% of the CAV-SCN1A-injected DS mice showed complete protection from seizures up to 40.5°C (**Fig. 3E**). Moreover, CAV-SCN1A injections in DS mice that exhibited seizures also demonstrated reduced susceptibility and an elevated temperature threshold (**Fig. 3E**). Together, these data demonstrate the potential of CAV-SCN1A treatment to improve the epileptic phenotypes in DS *Scn1a*^A1783V/WT^ mice during the chronic stage.

### Exogenous Na_V_1.1 activity does not alter the behavior of WT mice

Next, we compared the behavior of mock and treated adolescent WT and DS mice in the Y maze spontaneous alternation test and the open field test. The first uses the natural curiosity of mice and their tendency to explore novel environments to assess working memory, while the second monitors activity in a novel environment. While we did not detect significant differences between the groups (**Fig. 3F**), the alteration level in WT mice, injected with either CAV-GFP or CAV-SCN1A, was above chance level, indicating non-random exploration. Conversely, DS mice injected with CAV-GFP demonstrated random exploration (**Fig. 3F**), consistent with deficit in their working memory, while DS mice injected with CAV-SCN1A showed a tendency for more directed exploration, but with no clear evidence for correction (**Fig. 3F**). Interestingly, while CAV-SCN1A injections in the hippocampi had no adverse effects on the performance of WT mice in the open field test, hippocampal injections also did not rectify the hyperactivity of DS mice (**Fig. 3F**).

Together, these data indicate the lack of adverse effects of CAV-2 vectors, and exogenous Na_V_1.1 activity in the CNS of WT mice, while demonstrating the therapeutic potential for DS phenotypes when administered during the chronic phase in DS mice.

### CAV-SCN1A hippocampal injections revert epileptic phenotypes in juvenile DS mice

The severity of the epileptic phenotypes subsides after the 4^th^ week of life in DS *Scn1a*^A1783V/WT^ mice. Thus, we next asked if exogenous Na_V_1.1 activity, mediated by CAV-SCN1A injections, can prevent recurrent spontaneous seizures during the severe stage of the disease (38, 45). For this purpose, mice were administered bilateral hippocampal injections at the onset of symptoms (P21-P24). Of note, in juvenile mice, CAV-2-mediated GFP expression was also robust, with transduced neurons throughout the layers of the hippocampus and in hippocampal projecting regions including the neocortex and the thalamus (**Fig. S5**).

Examination of its therapeutic effect demonstrated that CAV-SCN1A injections significantly (*p* = 0.034) reduced SUDEP by ∼40% compared to mice injected with CAV-GFP (**Fig. 4A**). Continuous video recordings over 24 to 36 h post-injection of a subset of CAV-SCN1A-injected *Scn1a*^A1783V/WT^ mice, demonstrated that all the individuals experienced spontaneous convulsive seizures 12 h post-injection. Consistent with the rapid onset of transgene expression mediated by adenovirus vectors, by 36 h post-injection we found a significant (*p* < 0.05) reduction in the number of convulsive seizures in CAV-SCN1A-injected *Scn1a*^A1783V/WT^ mice, compared to DS mice injected with CAV-GFP (**Fig. 4B** **and Fig. S6**). CAV-SCN1A injections in juvenile DS mice either completely prevented thermally-induced seizures (∼30% of the treated mice) (**Fig. 4C**), or increased the seizure threshold temperature in mice that retained some sensitivity to thermally-induced seizures (**Fig. 4C**). Additionally, analyses of hippocampal and cortical recordings, performed in a subset of mice 2 weeks post-injection, revealed that CAV-SCN1A injections reduced the number of epileptic spikes observed in both regions (**Fig. 4D-G**). Together, these data demonstrate that CAV-mediated Na_V_1.1 activity, during the severe stage of DS, improved survival, reduced spontaneous seizures and epileptic spikes occurrence, and increased the temperature threshold of thermally-induced seizures.

**Fig. 4.**
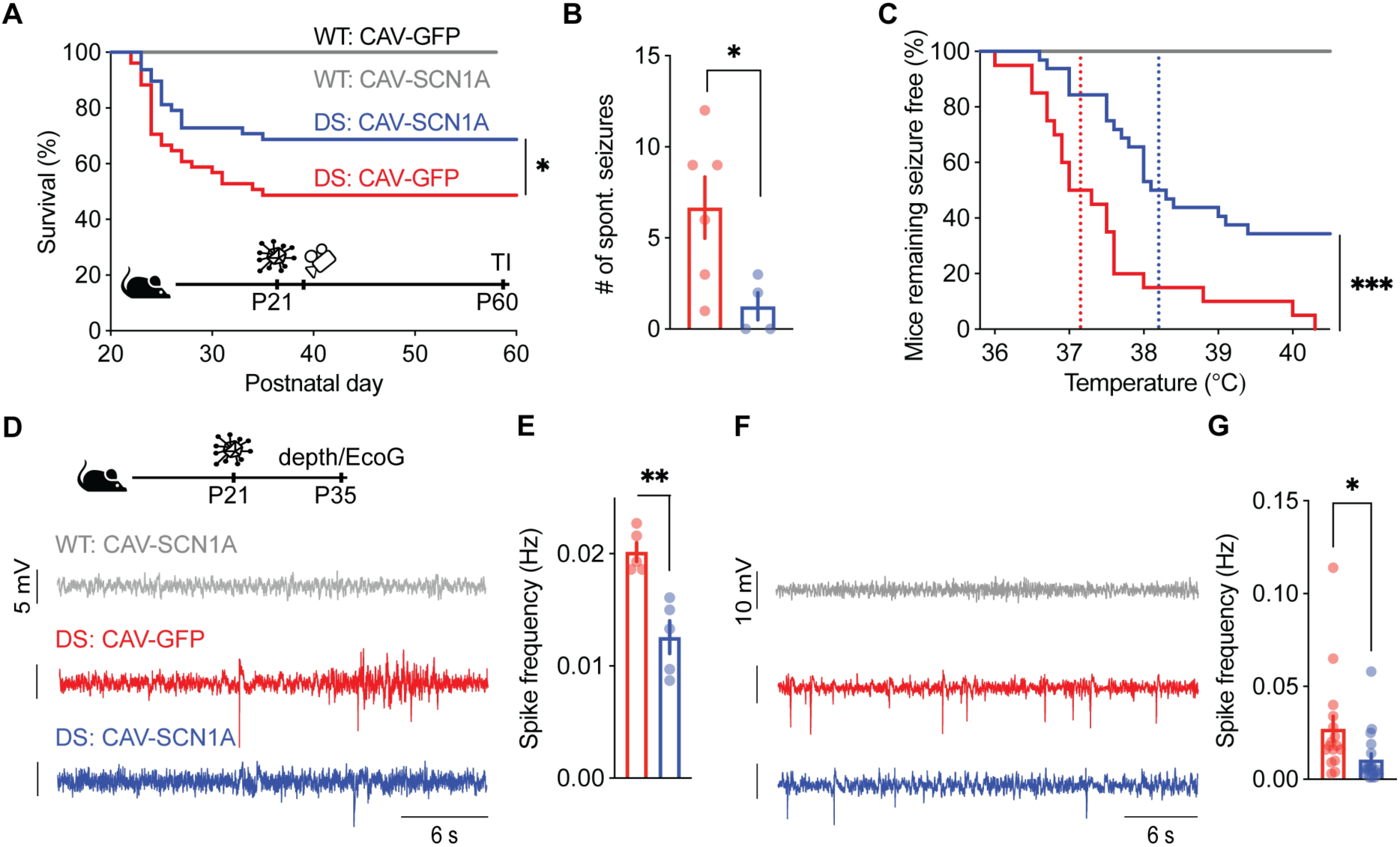
Hippocampal injection of CAV-SCN1A during the severe stage of Dravet ameliorates the epileptic phenotypes. (**A**) Survival curve of WT and DS littermates injected with either CAV-GFP or CAV-SCN1A at P21-P24. WT: CAV-GFP (n = 17); WT: CAV-SCN1A (n = 17); DS: CAV-GFP (n = 51); DS: CAV-SCN1A (n = 48). For separated analyses of males and females, see **Fig. S3**. (**B**) Video monitoring of convulsive seizures 36 h post-injection. DS: CAV-GFP (n = 6); DS: CAV-SCN1A (n = 4). See **Fig. S6** for additional data on individual mice. (**C**) Mice remaining free of thermally-induced seizures. The dotted lines represent the median seizure temperature. DS: CAV-GFP (n = 20); DS: CAV-SCN1A (n = 32). For separated analyses of males and females, see. **Fig. S3**. (**D-G**) Two weeks after the injections, depth electrodes (**D, E**) or cortical electrodes (**F, G**) were implanted. Example traces (**D, F**) and quantification of the spike frequencies are presented (**E, G**). DS: CAV-GFP (n = 5 for E, n = 16 for G); DS: CAV-SCN1A (n = 5 for E, n = 19 for G). No epileptic activity was detected in WT mice injected with either CAV-GFP or CAV-SCN1A (n = 5 each for depth electrodes and n = 9, n = 10 respectively, for cortical electrodes). See **Table S1** for details about statistical analyses.

### CAV-SCN1A injections restore background ECoG activity and partially correct behavioral deficits in DS mice

To further explore the impact of exogenous Na_V_1.1 activity, as well as the potential therapeutic effect of this treatment on background, non-epileptic brain oscillations, we performed spectral analysis of the ECoG signals approximately 2 weeks post-injection. The results indicate that injection of CAV-GFP or CAV-SCN1A did not alter the spectral ECoG profile of WT mice (**Fig. 5A-F**; data for untreated mice are replotted from (38)). We therefore concluded that neither CAV-2 vectors nor exogenous Na_V_1.1 activity, in excitatory and inhibitory neurons, impact global brain oscillations in a healthy brain.

**Fig. 5.**
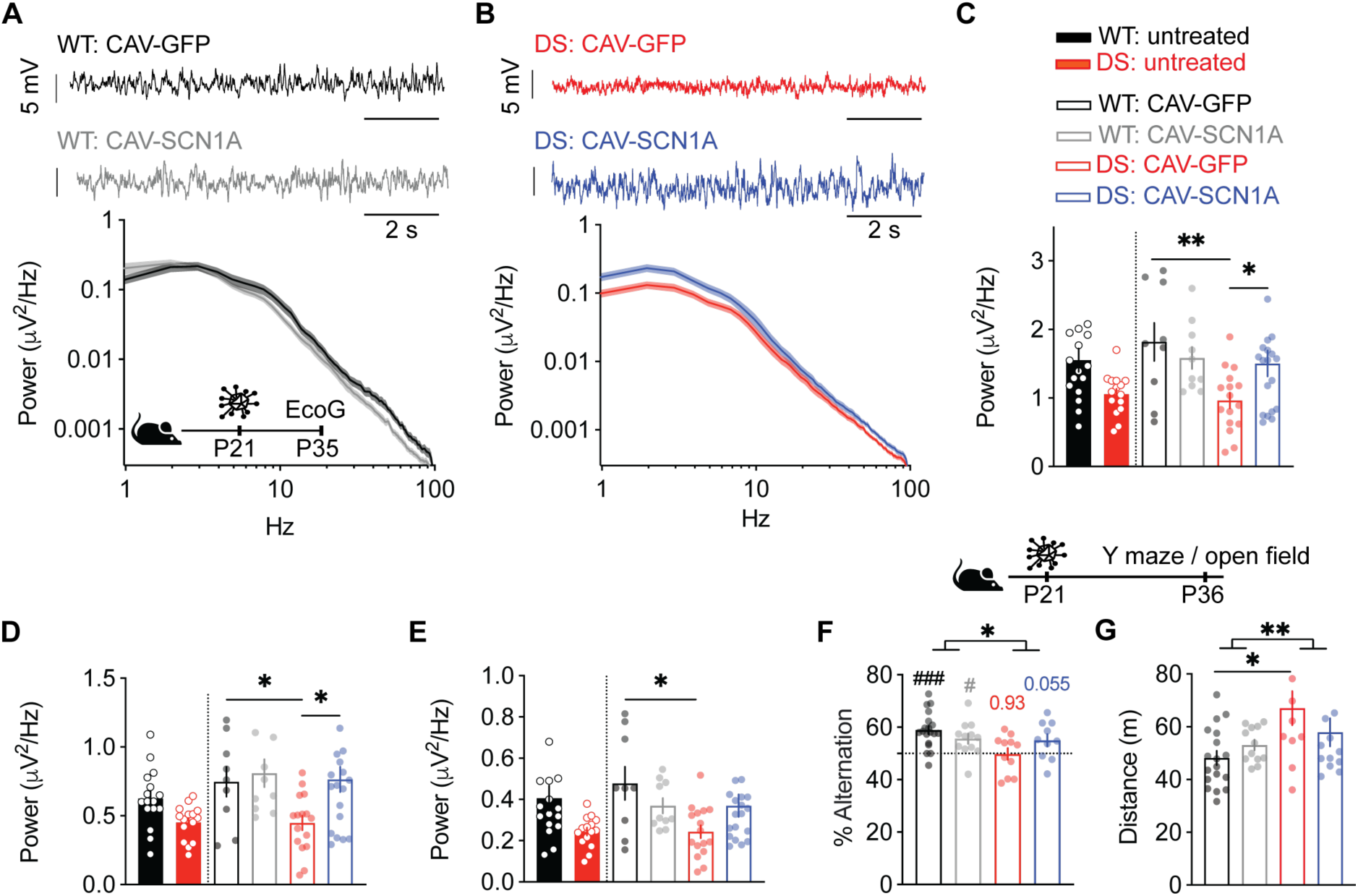
Correction of background ECoG activity, and partial correction of cognitive functions following CAV-SCN1A injections in juvenile DS mice. (**A-B**) Examples of background ECoG traces and power density profile of WT (**A**) and DS (**B**) mice. (**C-E**) Total power (**C**, 0.5-100 Hz), the power in the delta (**D**, 0.5-3.9 Hz) and theta bands (E, 4-8 Hz). Data for untreated mice are replotted from (Fadila et al., 2020). WT: CAV-GFP (n = 9); WT: CAV-SCN1A (n = 10); DS: CAV-GFP (n = 16); DS: CAV-SCN1A (n = 19). (F) Spontaneous alternation in the Y maze. The dotted line signifies the chance level, expected from random alternation. The markings above the bars indicate statistical analysis using one-sample *t* test relative to 50%. WT: CAV-GFP (n = 19); WT: CAV-SCN1A (n = 12); DS: CAV-GFP (n = 11); DS: CAV-SCN1A (n = 10). See Fig. S4, for separated analyses for males and females. (**G**) The distance moved in the open field. WT: CAV-GFP (n = 18); WT: CAV2-SCN1A (n = 12); DS: CAV-GFP (n = 11); DS: CAV-SCN1A (n = 13). See Fig. S4, for separated analyses for males and females. Statistical analysis between genotypes and treatments utilized Two Way ANOVA, See Table S1 for more details.

We have previously demonstrated that DS mice generally have lower power of background ECoG activity compared to WT mice (38). Accordingly, the power of the ECoG signals of DS mice injected with CAV-GFP, was lower compared to that of WT mice (*p* = 0.003), although similar to the power of untreated DS mice (**Fig. 5C**, data for untreated mice are replotted from (38)). Conversely, treatment with CAV-SCN1A resulted in higher power, particularly in the delta and theta frequency bands (**Fig. 5B-E**). This indicates that exogenous Na_V_1.1 activity generated by CAV-SCN1A injections also restores the low background ECoG activity in DS mice to WT levels. Because of the possibility of a physiological balance between the activity of sodium channels (9, 35, 49, 50), we also assessed if CAV-2 vector injections or exogenous Na_V_1.1 activity impacted the expression of endogenous *Scn1a, Scn2a, Scn3a, Scn8a* or the beta subunits*, Scn1b, Scn2b* and *Scn3b*. The results demonstrated that CAV-SCN1A injections did not alter the mRNA level of any of the voltage-gated sodium channels subtypes in control or DS mice (**Fig. S7**).

With respect to the effect on cognitive abilities, CAV-SCN1A injections had no effect on the performance of WT mice. However, there was a tendency (*p* = 0.055) for less random exploration in CAV-SCN1A-injected DS mice (**Fig. 5F**). Moreover, while CAV-GFP-injected DS mice traveled greater distances in the open field compared to WT mice treated with the same vector (**Fig. 5G**), CAV-SCN1A-injected DS mice traveled a similar distance to that of WT mice (**Fig. 5G**). We concluded that, in addition to the impact on epilepsy, CAV-SCN1A injections impacted non-epileptic DS features.

### Thalamic delivery of CAV-SCN1A revert epileptic phenotypes in juvenile DS mice

In addition to the impact on the hippocampus, thalamic dysfunction contributes to network hypersynchrony and DS (51–55). Therefore, we asked if thalamic injections of CAV-SCN1A could influence epilepsy in juvenile DS mice. Again, we found that CAV-SCN1A injections reduced the occurrence of SUDEP (**Fig. 6A**) and reduced the susceptibility to thermally-induced seizures (**Fig. 6B**). Furthermore, cortical ECoG recordings demonstrated that thalamic CAV-SCN1A injection can reduce the frequency of epileptic spikes (**Fig. 6C, D**) and correct the power of background activity (**Fig. 6E**). Together, these data demonstrate improvement of epileptic phenotypes and global brain oscillations following CAV-SCN1A injection into the thalamus of DS mice.

**Fig. 6.**
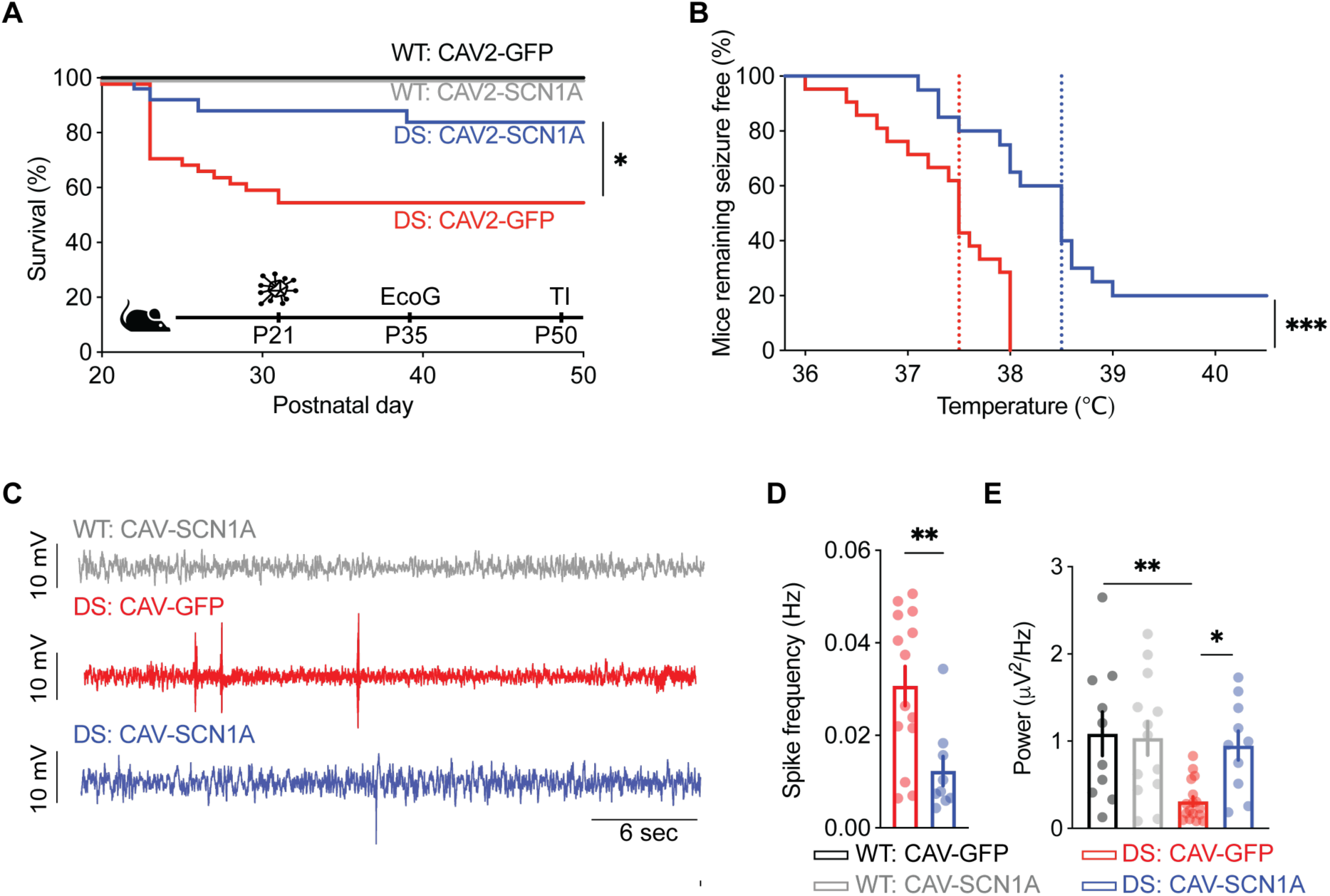
Thalamic injection of CAV-SCN1A ameliorates DS phenotypes in juvenile mice. (**A**) Survival curve of WT and DS littermates injected with either CAV-GFP or CAV-SCN1A. WT: CAV-GFP (n = 24); WT: CAV-SCN1A (n = 31); DS: CAV-GFP (n = 44); DS: CAV-SCN1A (n = 25). For separated analyses of males and females see **Fig. S8**. (**B**) Mice remaining free of thermally-induced seizures. The dotted lines represent the median seizure temperature. DS: CAV-GFP (n = 28); DS: CAV-SCN1A (n = 20). For separated analyses of males and females see **Fig. S8**. (**C-D**) Two weeks after the treatments cortical electrodes were implanted. Example traces (**C**) and quantification of the spike frequencies are depicted (**D**). (**E**) Total ECoG power (0.5-100 Hz) WT: CAV-GFP (n = 11); WT: CAV-SCN1A (n = 13); DS: CAV-GFP (n = 14); DS: CAV-SCN1A (n = 10). See **Table S1** for more details.

Following these results, we examined whether co-injections of CAV-SCN1A into the thalamus and hippocampus could further enhance the therapeutic effect of exogenous Na_V_1.1 activity. Indeed, following dual deposits along a unique needle track, DS mice treated with CAV-SCN1A demonstrated a significant (*p* = 0.007 and over 80%) increase in survival (**Fig. 7A**), as well as notable protection from thermally-induced seizures (**Fig. 7B**). Moreover, similar to the effect following hippocampal or thalamic administration, co-injections of CAV-SCN1A reduced the number of epileptic spikes and tended to increase the power of background ECoG (**Fig. 7C, D**). Therefore, treatment of juvenile DS mice with CAV-SCN1A in the hippocampus or the thalamus reduces their epileptic phenotypes, while combined delivery provides greater protection from thermally-induced seizures (**Fig. S10**).

**Fig. 7.**
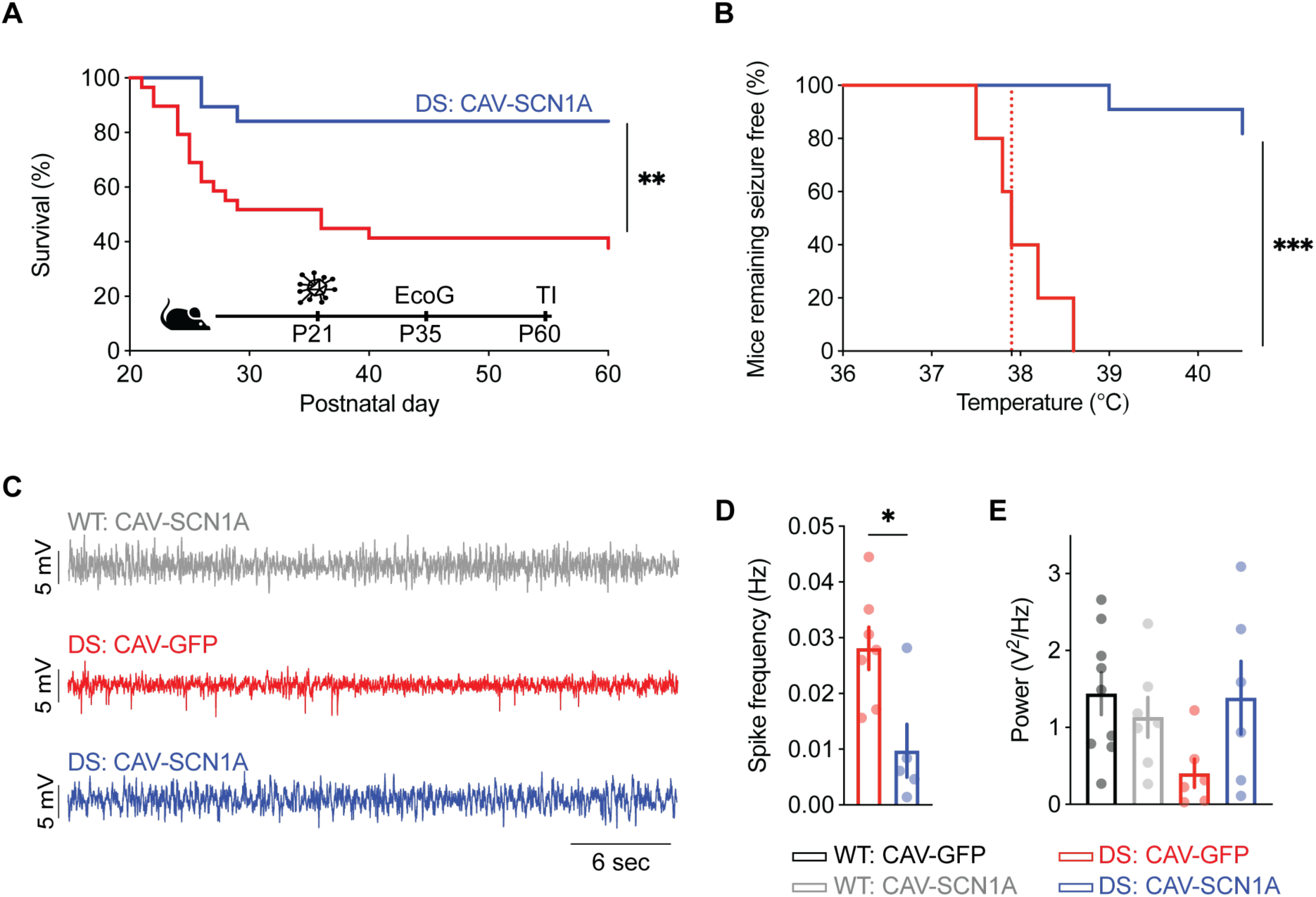
Concomitant thalamic and hippocampal injection of CAV-SCN1A in juvenile mice protects from thermally-induced seizures. (**A**) Survival of DS mice injected with either CAV-GFP or CAV-SCN1A. DS: CAV-GFP (n = 30); DS: CAV-SCN1A (n = 18). For separated analyses of males and females see **Fig. S9**. (**B**) Mice remaining free of thermally-induced seizures. The dotted lines represent the median seizure temperature. DS: CAV-GFP (n = 5); DS: CAV-SCN1A (n = 11). For separated analyses of males and females see **Fig. S9**. (**C-D**) Cortical electrodes were implanted two weeks after the treatments. Example traces (**C**) and quantification of the spike frequencies are depicted (**D**). (**E**) Total ECoG power (0.5-100 Hz) WT: CAV-GFP (n = 5); WT: CAV-SCN1A (n = 4); DS: CAV-GFP (n = 6); DS: CAV-SCN1A (n = 10). See **Table S1** for more details.

## Discussion

Dravet syndrome is an intractable childhood DEE, with a high fatality rate compared with other developmental epilepsies. SUDEP is the leading cause of death, with most fatalities occurring before the age of 10 (56). Importantly, despite polytherapy and recent advancements in therapeutic option, pharmacological seizure control in DS remains notoriously difficult (57). Therefore, there is an urgent need for novel treatments. In addition to the life-threatening epileptic seizures, individuals with DS suffer from non-epileptic comorbidities, including developmental delays, cognitive impairment, and hyperactivity. Although these behavioral deficits greatly impact the quality of life of patients and families, the therapeutic toolbox for addressing these burdensome issues is also limited. Here, we demonstrate that CAV-2-mediated exogenous Na_V_1.1 activity, via a codon-modified *SCN1A* cDNA, can significantly improve comorbidities in juvenile and adult DS mice, ameliorate the epileptic phenotypes, correct background ECoG activity, and tend to improve cognitive functions. Notably, this potent treatment can be adapted for clinical use during the severe or the chronic stages of DS.

C57BL/6J-*Scn1a*^A1783V/WT^ mice display particularly severe epileptic phenotypes compared to other DS mice (7, 8), which may render this model more resistant to treatment. Specifically, like human patients with DS, C57BL/6J-*Scn1a*^A1783V/WT^ mice experience spontaneous seizures (**Fig. S6** and (45)) and thermally-induced seizures that occur within the range of clinical febrile seizures (58). Furthermore, while the frequency of SUDEP is higher than the risk reported in patients (**Figs. 4, 6, 7**) (38, 59), it must be appreciated that the mice do not receive anti-seizure medication or emergency care for prolonged seizures. Thus, we argue that the severe epileptic phenotype of C57BL/6J-*Scn1a*^A1783V/WT^ mice faithfully represents the clinical severity of DS. Importantly, in contrast to other models that harbor *Scn1a* truncation mutations, effective gene therapy in this model needs to overcome the endogenous membrane location of Na_V_1.1 channels with reduced activity (41). Inasmuch, the therapeutic impact of CAV-SCN1A in this challenging DS model, highlights the clinical potential of this approach.

Notably, CAV-SCN1A demonstrated similar efficacy at multiple disease stages (**Figs. 3-7**). This is in contrast to previous attempts to provide gene-specific treatments for DS, which demonstrated a therapeutic effect when administered during the asymptomatic, pre-epileptic stage (5, 7, 8), or during the chronic phase (6, 29). When administered during the severe stage of the disease, CAV-SCN1A improved the survival, reduced the occurrence of spontaneous seizures, the frequency of epileptic spikes and protected from thermally-induced seizures (**Figs. 4, 6, 7**). Because the frequency of spontaneous seizures and occurrence of SUDEP is low during the chronic stage of the disease, therapeutic efficacy was demonstrated by reduced frequency of epileptic spikes and reduced sensitivity to thermally-induced seizures (**Fig. 3**). Moreover, as our treatment relies on vector-mediated *SCN1A* cDNA delivery, rather than transcriptional activation of a functional (and mutated) *SCN1A* alleles, it is suitable for patients with truncation or missense mutations in *SCN1A*.

Despite our advances, there was still some residual epileptic activity after therapy (**Figs. 3-7**). This is in accordance with reports that spontaneous seizures were still observed when antisense oligonucleotides designed to enhance *Scn1a* transcription were administered at P2, well within the asymptomatic period, and did prevent premature death. Moreover, antisense oligonucleotide treatment given at P14, still in the midst of the pre-epileptic stage, could not achieve full protection from death (7). Similarly, epileptic spikes and spontaneous seizures were still detected following neonate dCas9-based *Scn1a* activation (5), and SUDEP (despite improved survival) was still evident following vector-mediated overexpression of Na_V_β1 (8). Likewise, in most cases abnormal epileptic activity still persists when the treatments were administered during the chronic stage of the disease (6, 29).

In our model, administration of CAV-SCN1A to the hippocampus or the thalamus, had a similar effect on DS epilepsy (**Figs. 4, 6**), consistent with an involvement of these areas in DS pathophysiology. Concomitant delivery into these two sites afforded similar protection from SUDEP (**Fig. S10A**), but provided greater protection from thermally-induced seizures (**Fig. S10B**), possibly due to a wider biodistribution of the exogenous functional Na_V_1.1. Interestingly, distinct circuit-specific neuronal dysfunctions have been described for each of these regions. Disinhibition was proposed as the culprit in the hippocampus (21, 25, 35, 37, 43, 45, 60). Conversely, complex neuronal changes were reported in the thalamus, with reduced activity of inhibitory and excitatory neurons (52, 53, 55), as well as hyperexcitability of inhibitory thalamic reticular nucleus neurons that led to augmented cortico-thalamic oscillations and seizures (51). Thus, the therapeutic effect of CAV-SCN1A in both locations, despite the seemingly opposing local neuronal dysfunctions, further emphasize the potential of our approach.

One significant challenge for CNS-targeted gene delivery is the need to transduce enough neurons within a critical brain region, to trigger a global change in network function. Our data demonstrate the pivotal role of Na_V_1.1 expression within the hippocampus and thalamus for DS therapy. Within the hippocampus, HA-SCN1A immunoreactivity was mostly detected in inhibitory neurons (**Fig. 2**), in agreement with the contribution of these cells to DS pathophysiology (19, 20, 39), and preferential expression of Na_V_1.1 in inhibitory neurons (33, 61). Thus, we propose that CAV-SCN1A transduction of a small number of critical neurons may be sufficient for effective therapy. In support of this notion, it was recently shown that disruption of the activity of single neurons, within a connected hub can disrupt epileptic network dynamics (62).

Nevertheless, the NSE promoter does not limit the expression to inhibitory neurons (**Figs. 1, S5**). Therefore, we cannot exclude that possibility that restoration of low levels of Na_V_1.1 expression in excitatory neurons contributes to the observed effects. Indeed, one of the advantages of CAV-2 vectors, is their retrograde transport to neurons projecting into the structure injected, leading to widespread CNS gene expression (13, 14, 17). Thus, it is possible that providing exogenous Na_V_1.1 activity, within a tightly connected neuronal network that hubs at the injection site, but includes multiple projecting neurons (mostly excitatory) from various additional brain regions (**Figs. 1, 2, S5**), contributes to the observed global effect.

As with most initial attempts, there remains room for certain improvements of the therapeutic potential of exogenous Na_V_1.1 activity mediated by CAV-2 vectors. Near global brain reactivation of *Scn1a* expression in the mouse brain provided an encouraging roadmap towards DS therapy (10). However, genetically modifying all the neurons in the human brain will never be possible -or needed - for DS therapy. Here is where CAV-2 vectors can have an additional impact on the fundamental understanding of DS inception and progression. Using transcriptional and/or translation control elements we can target Na_V_1.1 activity to specific neuronal subpopulations, in specific regions of the brain, at a given age, to identify their role during the evolution of DS. Then, these data will eventually allow us to further improve CAV-2-mediated *SCN1A* cDNA transfer to treat all DS comorbidities. There are several additional avenues to explore including optimizing the dose injected (NB: we injected only 10^9^ physical particles), and/or the inclusion of additional expression cassettes in HD CAV-2 vectors that could impact epileptic activity. Moreover, exogenous Na_V_1.1 activity may synergize with pharmacological approaches to further improve DS therapy.

In conclusion, these results provide a proof-of-concept for the potential of CAV-mediated SCN1A delivery as a therapeutic approach for children and adolescents with DS-associated *SCN1A* missense and truncation mutations.

## Methods

### Vectors generation

All CAV-2 vectors used in this work were generated using a seamless ligation cloning extract (SLiCE) strategy (63). The following promoters were obtained from Addgene: the NSE promoter (plasmid # 50958 James Bamburg, (64)); the CAG promoter (plasmid # 51274, Pawel Pelczar, (65)) ;hSyn promoter (plasmid # 22909, Edward Callaway, (66)); the Dlx5/6 enhancer (plasmid # 83900, Gordon Fishell, (28)), mCitrine, SCN1A cDNA (29) and the bovine growth hormone polyA sequence. Fragments were subcloned into the E1 region of E1/E3-deleted pCAV-2. In order to avoid vector genome rearrangements/deletions during amplification (∼50,000 vector genomes are produced /cell), we modified the *SCN1A* cDNA using codon optimization algorithms. This automated step was followed by manually screening and further modification of short (10-15 bp) repeat sequences mainly in the transmembrane coding regions. Vectors were expanded and purified as previously described (30, 67).

### Animals

Animal handling was conducted in accordance with the European Council directive (2010/63/EU) and the ARRIVE guidelines, and approved by appropriate Intuitional Ethical Committee. WT and DS mice harboring the global *Scn1a*^A1783V/WT^ mutation were generated by crossing conditional floxed *Scn1a*^A1783V^ males (The Jackson Laboratory; stock #026133) with CMV-Cre females (stock #006054). Mice were housed in a standard animal facility at a constant temperature of 22°C, on a 12-hour light/dark cycle, with *ad libitium* access to food and water.

### Vector injections

Adult (8-14 weeks) WT C57BL/6J mice were anesthetized with 90 mg/kg ketamine,10 mg/kg xylazine and 2 mg/kg of acepromazine by intraperitoneal injections resulting in deep anaesthesia. The animal was positioned in the stereotaxic frame; the coordinates for the hippocampus were: anteriorposterior (AP) -2.1 mm; mediolateral (ML) ±1.5 mm; dorsoventral (DV) -1.8 mm (68), 2 μl containing 1×10^9^ physical particles were injected on each side, using a 5 μl Hamilton syringe (0.2 μl/min).

WT and DS mice at the age of P21-24, or P35-P36 were randomly assigned to treatment with CAV-2-NSE-GFP (CAV-GFP) or CAV-2-NSE-SCN1A (CAV-SCN1A). The mice were anesthetized using ketamine/xylazine (191/4.25 mg/kg), carprofen (5 mg/kg) was used for analgesia. The mice were placed in a stereotaxic device (Ultra Precise Stereotaxic Instruments, Stoelting, Wood Dale, IL, USA). A midline incision was made above the skull and holes were made using a 25G needle at the site of injection. For hippocampal injections we used the following coordinates: AP -1.8 mm; ML ±1.7 mm; DV -3 mm for P21-24 and AP -1.7 mm; ML ±1.8 mm; DV -3 mm for P35-36. For thalamic injections at P21-P24 we use the following coordinates: AP -1.4 mm; ML ±2.2 mm; DV -3.5 mm. 1 μl containing 1×10^9^ physical particles were injected on each side, using a 1 μl beveled needle Hamilton syringe (1 μl syringe, 7000 series, beveled tip).

For dual thalamic-hippocampal injections the injection needle was first lowered to: AP -1.8 mm; ML ±1.8 mm; DV -3.5 mm, and a volume of 0.5 μl was injected. After a few min, the injection needle was raised to DV 3 mm, and another volume of 0.5 μl was injected. The AP coordinates were slightly modified for P21 mice in which the distance between bregma and lambda was less than 3 mm, and reduced to AP -1.7 mm. The DV coordination was measured from the tip of the beveled injection needle. CAV vectors were injected at a rate of 100 nL/min (Quintessential Stereotaxic Injector, Stoelting, Wood Dale, IL, USA). After injection, the syringe was kept in place for at least 5 minutes to prevent backflow before it was slowly retracted. The skin was then closed with sutures and the mice were isolated for a period of 7 days according to TAU BSL-2 safety instructions.

### Immunohistochemistry

Ten-days post-surgery, the animals were injected with an overdose with ketamine and xylazine and perfused transcardially with saline buffer (0.9%) followed by a fixative solution containing 4% paraformaldehyde (PFA) and phosphate buffer (PB) 0.1M, pH 7.4. The brains were post-fixed in 4% PFA for 24 h and soaked in 30% sucrose prepared in PB 0.1M at 4°C for at least 48 h. Next, the brains were frozen in Tissue Freezing Medium® optimum cutting temperature (O.C.T.) (MicromMicrotech, TFM-5) and stored at -80°C. OCT blocks were cut in serial coronal sections (35 μm thick) using a cryostat and collected in a solution containing glycerol and ethylene glycol in PB.

IH on Free-floating sections was performed as previously described (16). The following primary antibodies were used: chicken anti-GFP (1:1000, ABCAM, ab13970 RRID:AB_300798), a rat anti-HA (1:200 Roche, 11 867 431 001) anti-GABA (Sigma-Aldrich, SAB4200721, RRID:AB_2891218). Following the incubation with primary antibodies, sections were rinsed with TBST and incubated with the appropriated biotinylated or fluorescent secondary antibody/antibodies diluted in the blocking solution for 1 h. The following secondary antibodies were used in this study: donkey anti-chicken (1:500, Jackson ImmunoResearch Labs, Cat# 703-065-155, RRID:AB_2313596), 4′,6-diamidino-2-phenylindole dihydrochloride (Sigma D8417). The sections processed for colorimetric immunohistochemistry were incubated with avidin-biotin-complex (Vector Laboratories PK-6100, RRID:AB_2336819) for 1 h at room temperature. Once washed in TBS, the peroxidase reaction was visualized using 0.05% 3,3′-diaminobenzidine (Sigma, D5637) and 0.03% hydrogen peroxide. Finally, sections were rinsed in TBS and mounted on SuperFrost Ultra Plus® slides, dried at room temperature and counterstained using Cresyl violet, dehydrate and coverslipped with Eukitt (Sigma, 03989).

To test the specificity of the secondary antibodies, we omitted the primary antibodies in some sections while maintaining the rest of the procedures. All the control sections exhibited a lack of positive staining.

The colorimetric signals were visualized using a NIKON ECLIPSE NI-E and a color camera DS-Ri2 (4908*3264 px de 7.3 μm) or Nanozoomer (Hamamatsu). Images were adjusted for brightness and contrast by using ImageJ. Picture setup was achieved with Adobe Illustrator CS6. Full resolution was maintained until the micrographs were cropped and assembled; at which time they were adjusted to a resolution of 300 dpi. The brain regions were identified using a mouse brain atlas (68).

### ECoG and depth electrode and recordings

Seven to ten days following vector injections, cortical or depth electrodes were implanted as previously described (38). Briefly, a midline incision was made above the skull, and fine silver wire electrodes (130 μm diameter bare; 180 μm diameter coated) were implanted. We used the previously formed injection holes for ECoG or depth electrodes. For hippocampal depth recordings, the wire electrodes were lowered using the same stereotactic coordinates used for injection. A reference electrode was placed on the cerebellum; a ground electrode was placed behind the neck. The electrodes were connected to a Mill-Max connector and secured with dental cement, before the skin was closed with sutures. Following the surgery, the mice were given two to five days to recover before recording.

Video-depth/EcoG recordings lasting 2-4 h were obtained during the light period from freely moving mice, connected to a T8 Headstage (Triangle BioSystems, Durham, NC, USA), using a PowerLab 8/35 acquisition hardware and the LabChart 8 software (ADInstrumnts, Sydney, Australia). The electrical signals were recorded and digitized at a sampling rate of 1 kHz with a notch filter at 50 Hz. The analysis was performed using LAbChart 8 (ADInstrumnts, Sydney, Australia). The ECoG signal was processed offline with a 0.5-100 Hz bandpass filter. Power spectral density was calculated using fast Fourier transform, with the Hann (cosine-bell) data window set to 50% overlap. We calculated the average of four to eight 30 s long segments of the wakefulness, immobility and epileptic-free activity, but following a movement for each mouse, as determined by the video recording.

### Thermally-induced seizures

Thermal induction was done about one month post injection as described previously (45). Briefly, the mice were given 10 minutes to habituate to the thermal probe and the recording chamber. The baseline body temperature was measured, followed by an increase of 0.5°C every 2 minutes until 40.5°C or until a seizure was generated.

### Behavioral experiments

Behavioral experiments were performed as described previously (38). Spontaneous alternation in the Y maze was assessed 5-14 days post-injection and the open field test was conducted 9-17 days after treatment. Briefly, for the Y-maze spontaneous alternation test the mice were placed in a symmetrical Y-maze composed of three opaque white Plexiglas arms (each 35 cm L x 7.6 cm W x 20 cm H) and allowed free exploration for 10 min. For the open-field test, mice were placed in the center of a square (50 x 50 cm) Plexiglas apparatus and their activity was recorded for 10 min. Live tracking was achieved via a monochrome camera (Basler acA1300-60gm, Basler AG, Ahren, Germany) connected with EthoVision XT 13 software (Noldus Technology, Wageningen, Netherlands).

### Statistics

Statistical analysis was performed using GraphPad Prism 9.2 (GraphPad Software, La Jolla, CA, USA), utilizing Log-rank test, Two-Way ANOVA, unpaired *t*-test or the Mann-Whitney test, as appropriate. The specific tests that were used for each panel are specified in **Table S1**.

## Study approval

All animal experiments were approved by the Ethical Committee for Animal Testing (Comité régional Languedoc-Roussillon) and the institutional care and use committee of Tel Aviv University.

## Author contributions

S.F, B.B, I.G.D.R designed and carried out the experiments, performed data analysis, and helped construct the manuscript. A.M, M.B, and K.A carried out some experiments and provided technical support. E.M.G, A.R, and R.H.A designed some experiments and provided critical analyses. E.J.K and M.R coordinated this study, secured funding, designed the experiments, and wrote the manuscript. All authors approved the final manuscript.

## Acknowledgments

We acknowledge the financial support of the E-Rare (to EJK, RH & MR). The Spanish Dravet Syndrome Foundation (RH), The American Dravet Syndrome Foundation (EJK, MR & EG), the ANR (DSynchro # ANR-21-CE17-0056-01, NORAD ANR-19-CE37-0008 (EJK)), the Israel Science Foundation (# 1454/17, MR), the Fondation pour la Recherche Médicale (EJK). We thank the members of the Rubinstein, Kremer and Hernandez labs for constructive comments during the course of the study. EJK is an Inserm Fellow.

