## Supplementary Materials: Figures S1-S10, Table S1 for "Exogenous Na_V_1.1 activity in excitatory and inhibitory neurons reverts Dravet syndrome comorbidities when delivered post-symptom onset in mice with Dravet"

| Fig | Panel | Test used | ***p* value and additional information** |
|---|---|---|---|
| Fig. 3 | B | unpaired t-test | *p* = 0.013 (*) |
| | D | unpaired t-test | *p* = 0.048 (*) |
| | E | Log-rank test | DS: CAV-GFP vs. DS: CAV-SCN1A<br>*p* < 0.001 (***) |
| | F<br>Y maze | One sample t-test | *p* values are marked on the graph |
| | | Two-way ANOVA | Main effect for genotype *p* = 0.078<br>Main effect for viral treatment *p* = 0.7<br>Genotype x viral treatment *p* = 0.72 |
| | F<br>Open field | Two-way ANOVA | Main effect for genotype *p* = 0.0009 (***)<br>Main effect for viral treatment *p* = 0.32<br>Genotype x viral treatment *p* = 0.31<br>Followed by Holm-Sidak post hoc analysis |
| Fig. 4 | A | Log-rank test | DS: CAV-GFP vs. DS: CAV-SCN1A<br>*p* = 0.034 (*) |
| | B | unpaired t-test | *p* = 0.038 (*) |
| | C | Log-rank test | DS: CAV-GFP vs. DS: CAV-SCN1A<br>*p* < 0.0001 (***) |
| | E | Mann-Whitney test | *p* = 0.0035 (**) |
| | F | unpaired t-test | *p* = 0.0019 (**) |
| Fig. 5 | C | Two-way ANOVA | Main effect for genotype *p* = 0.006 (**)<br>Main effect for viral treatment *p* = 0.48<br>Genotype x viral treatment *p* = 0.01 (*)<br>Followed by Holm-Sidak post hoc analysis |
| | D | Two-way ANOVA | Main effect for genotype *p* = 0.067<br>Main effect for viral treatment *p* = 0.87<br>Genotype x viral treatment *p* = 0.034 (*)<br>Followed by Holm-Sidak post hoc analysis |
| | E | Two-way ANOVA | Main effect for genotype *p* = 0.03 (*)<br>Main effect for viral treatment *p* = 0.15 |

| | | | |
|---|---|---|---|
| | | | Genotype x viral treatment $p = 0.037$ (*)<br>Followed by Holm-Sidak post hoc analysis |
| | F | One sample t-test | $p$ values are marked on the graph |
| | | Two-way ANOVA | Main effect for genotype $p = 0.016$ (*)<br>Main effect for viral treatment $p = 0.61$ Genotype x viral treatment $p = 0.036$ (*)<br>Followed by Holm-Sidak post hoc analysis |
| | G | Two-way ANOVA | Main effect for genotype $p = 0.007$ (**)<br>Main effect for viral treatment $p = 0.62$<br>Genotype x viral treatment $p = 0.1$<br>Followed by Holm-Sidak post hoc analysis |
| Fig 6 | A | Log-rank test | DS: CAV-GFP vs. DS: CAV-SCN1A<br>$p = 0.014$ (*) |
| | B | Log-rank test | DS: CAV-GFP vs. DS: CAV-SCN1A<br>$p < 0.0001$ (***) |
| | D | Mann-Whitney test | $p = 0.007$ (**) |
| | E | Two-way ANOVA | Main effect for genotype $p = 0.012$ (*)<br>Main effect for viral treatment $p = 0.08$<br>Genotype x viral treatment $p = 0.041$ (*)<br>Followed by Holm-Sidak post hoc analysis |
| Fig. 7 | A | Log-rank test | DS: CAV-GFP vs. DS: CAV-SCN1A<br>$p = 0.007$ (**) |
| | B | Log-rank test | DS: CAV-GFP vs. DS: CAV-SCN1A<br>$p < 0.0001$ (***) |
| | D | unpaired t-test | $p = 0.0002$ (***) |
| | E | Two-way ANOVA | Main effect for genotype $p = 0.22$<br>Main effect for viral treatment $p = 0.27$<br>Genotype x viral treatment $p = 0.097$<br>Followed by Holm-Sidak post hoc analysis |

**Table S1. Additional statistical information**

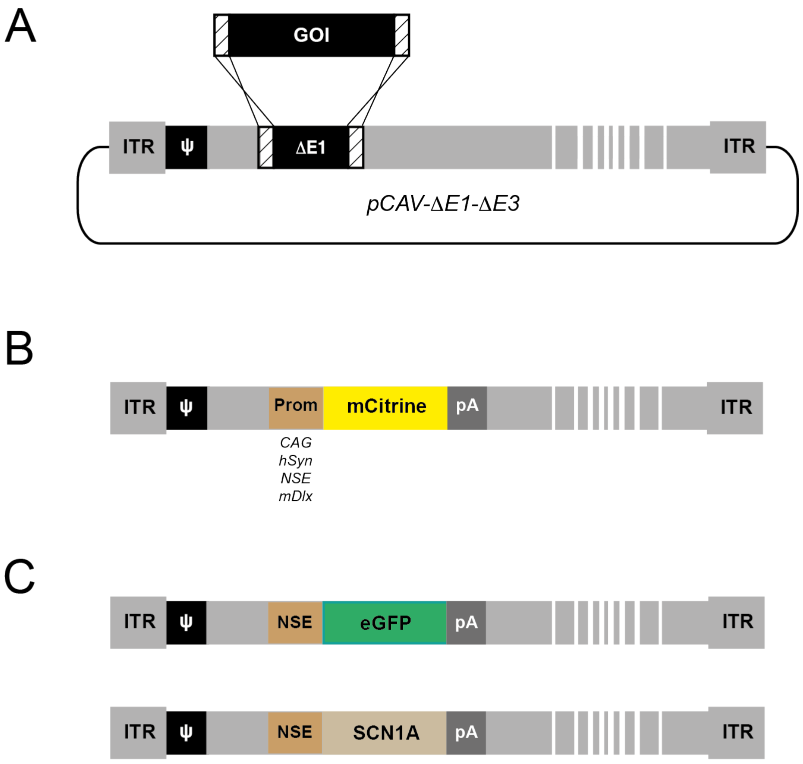


**Fig. S1. Description of CAV vectors used in this work.**

(**A**) Cloning strategy using SLiCE to insert cassette with gene of interest (GOI) in plasmid pCAV-ΔE1-ΔE3 carrying CAV genome deleted in E1 and E3 regions (in grey). ITR: inverted terminal repeat.

(**B**) Map of CAV vector used to study transcriptionally-targeted transgene expression (Prom = promoter ; pA = BGHpA). The promoters/enhancers were used: CAG, hSyn, NSE, mDlx.

(**C**) Map of CAV vectors used to study therapeutic effect *in vivo*. CAV carrying eGFP was used as control, and the vector harboring the codon-modified *SCN1A* cDNA as therapeutic vector.

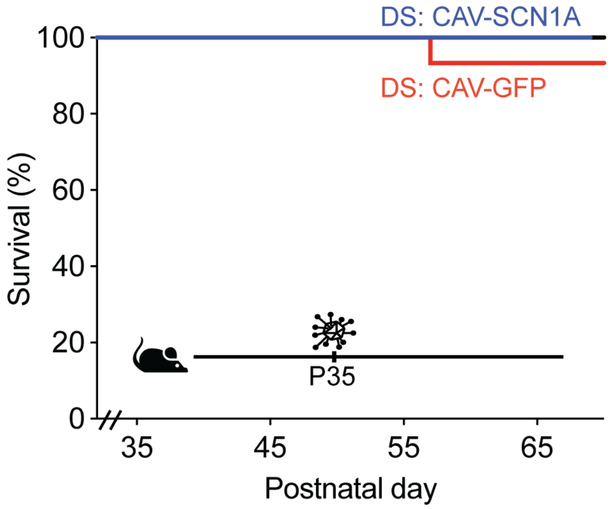


**Fig S2. Survival of adolescent mice following injection of CAV-GFP or CAV-SCN1A.**
Premature death in DS mice mostly occurs at the 4$^{th}$ week of life. Here we tested the ability of CAV-SCN1A to ameliorate the epileptic phenotypes in a subset of mice that survived to the 5$^{th}$ week of life. As expected at this age, only one DS mouse died prematurely (treated with CAV-GFP). WT: CAV-GFP n = 14; WT: CAV-SCN1A n =13; DS: CAV-GFP n = 15; DS: CAV-SCN1A n = 13.

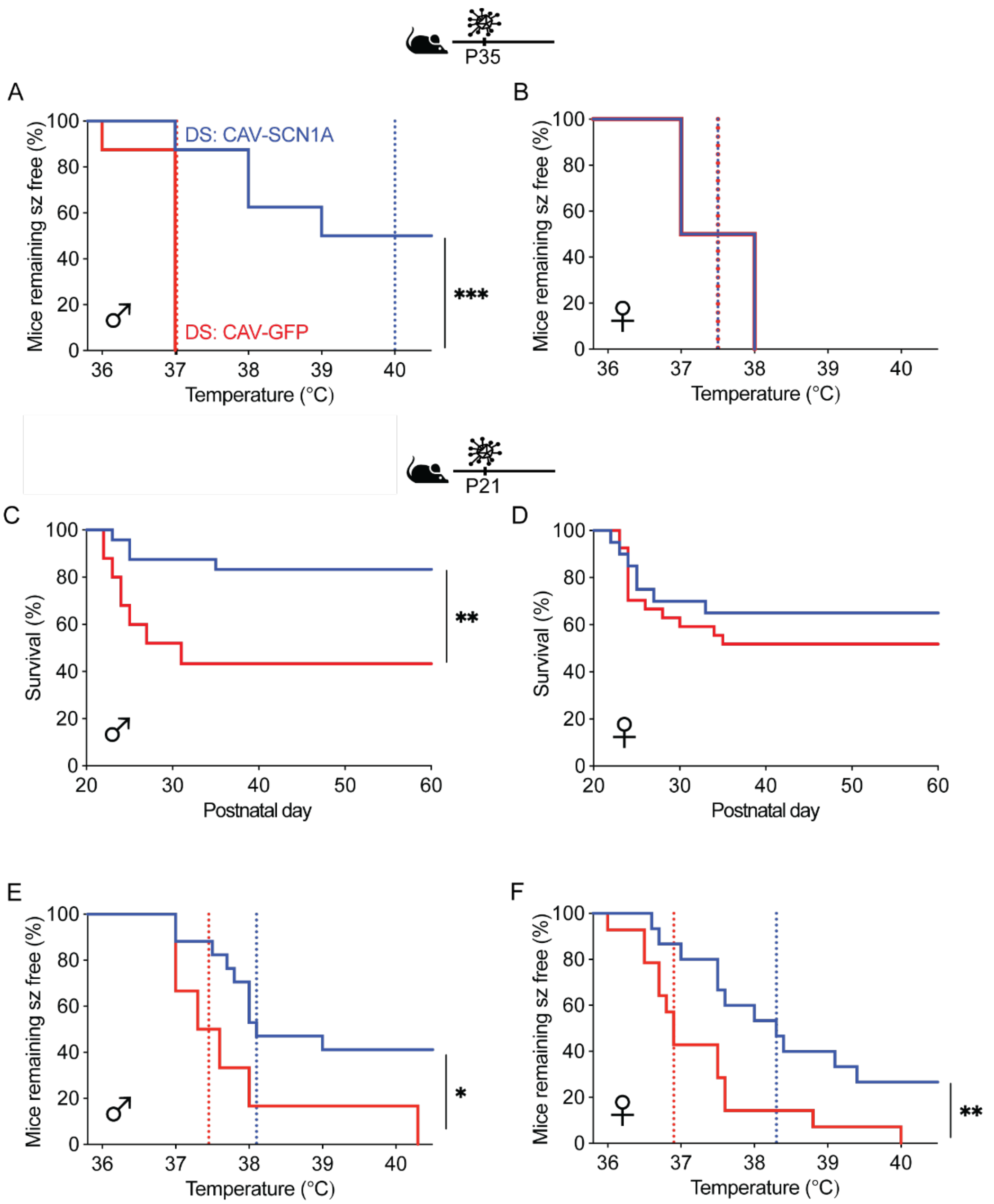


**Fig. S3. Some sexual divergent effects following CAV-SCN1A injections into the hippocampus.** (**A-B**) Males (**A**) and females (**B**) DS mice remaining free of thermally-induced seizures following hippocampal injections of CAV-GFP or CAV-SCN1A at 5 weeks of age. The dotted lines represent the median seizure temperature. DS: CAV-GFP, ♂ n = 8, ♀ n = 2; DS: CAV-SCN1A, ♂ n = 8, ♀ n = 6.
(**C-D**) Survival curve of males (**C**) and females (**D**) DS mice injected with either CAV-GFP or CAV-SCN1A at P21. DS: CAV-GFP, ♂ n = 25, ♀ n = 27; DS: CAV-SCN1A, ♂ n = 25, ♀ n = 20.
(**E-F**) Males (**E**) and females (**F**) DS mice remaining free of thermally-induced seizures. The dotted lines represent the median seizure temperature. DS: CAV-GFP, ♂ n = 6, ♀ n = 14; DS: CAV-SCN1A, ♂ n = 17, ♀ n = 15. Statistical analysis utilized the Log-rank test. *$p < 0.05$; ** $p < 0.01$; ***$p < 0.001$

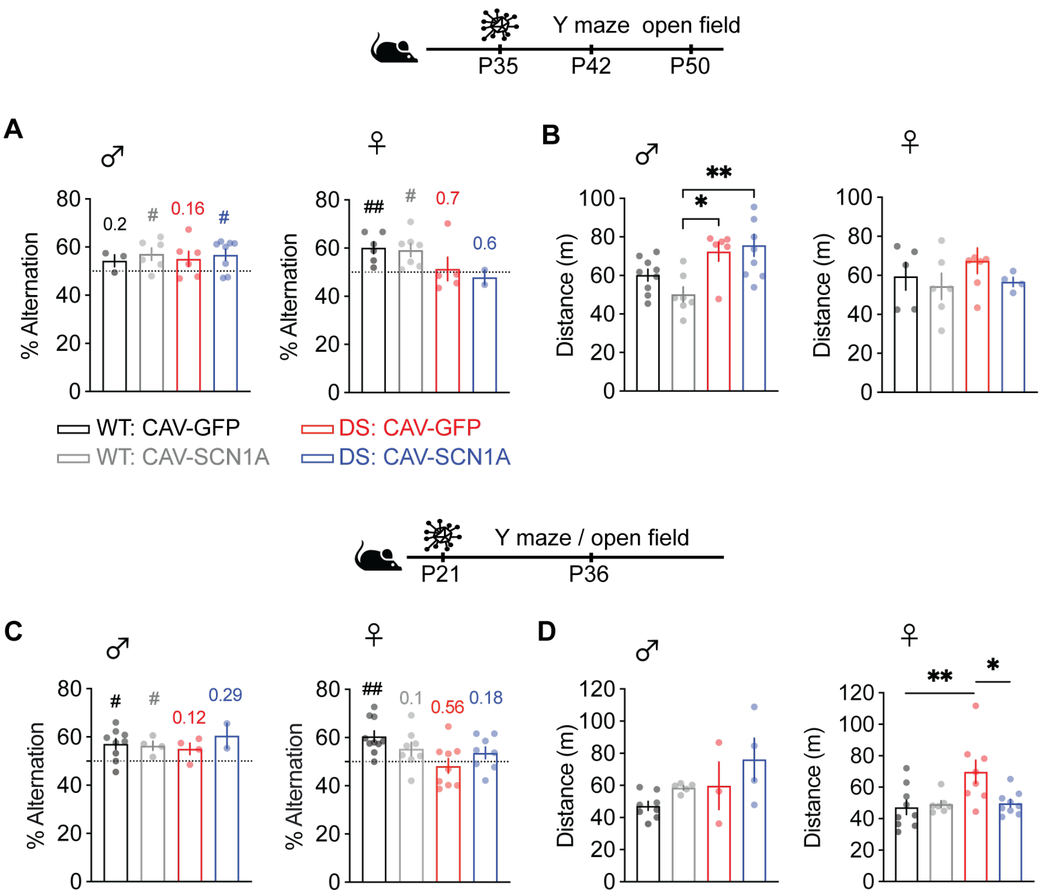


**Fig. S4. Behavioral characterization of male and female mice.**

(**A**) Percent spontaneous alternation in the Y maze, of males and females, following gene therapy in adolescent mice. The test was performed 5-10 days post injection (median age P42, range P39-P43). The dotted line signifies chance level, expected from random alternations. The markings above the bars indicate statistical analysis using one-sample *t* test relative to 50%. WT: CAV-GFP, ♂ n = 3; ♀ n = 6; WT: CAV-SCN1A, ♂ n = 6, ♀ n = 7; DS: CAV-GFP, ♂ n = 6, ♀ n = 5; DS: CAV-SCN1A, ♂ n = 8; ♀ n = 2.

(**B**) The open field test was performed 9-17 days post injection (median age P50, range P45-P53). WT: CAV-GFP, ♂ n = 9, ♀ n = 5; WT: CAV-SCN1A, ♂ n = 7, ♀ n = 6; DS: CAV-GFP, ♂ n = 6, ♀ n = 7; DS: CAV-SCN1A, ♂ n = 9, ♀ n = 4. Statistical analysis utilized two-way ANOVA: ♂ main effect for genotype $p$ = 0.0004 (***), main effect for viral treatment $p$ = 0.46, genotype x viral treatment $p$ = 0.16. The results of Holm-Sidak post hoc analysis are depicted on the graph. ♀ main effect for genotype $p$ = 0.45, main effect for viral treatment $p$ = 0.24, genotype x viral treatment $p$ = 0.67.

(**C**) Percent spontaneous alternation of males and females in the Y maze, following gene therapy in juvenile mice (P21). This test was performed 8-14 days post injection (median age P36, range P29-P39). The dotted line signifies chance level, expected from random alternations. The markings above the bars indicate statistical analysis using one-sample *t* test relative to 50%. WT: CAV-GFP, ♂ n = 9; ♀ n = 10; WT: CAV-SCN1A, ♂ n = 4, ♀ n = 8; DS: CAV-GFP, ♂ n = 4, ♀ n = 9; DS: CAV-SCN1A, ♂ n = 2, ♀ n = 8.

(**D**) The distance moved in the open field following gene therapy in juvenile mice. This test was performed 9-17 days post injection (median age P37, range P30-P41). WT: CAV-GFP, ♂ n = 8; ♀ n = 9; WT: CAV-SCN1A, ♂ n = 5, ♀ n = 7; DS: CAV-GFP, ♂ n = 3, ♀ n = 8; DS: CAV-SCN1A, ♂ n = 9, ♀ n = 4. Statistical analysis utilized two-way ANOVA: ♂ main effect for genotype $p = 0.06$, main effect for viral treatment $p = 0.08$, genotype x viral treatment $p = 0.73$. ♀ main effect for genotype $p = 0.019$ (*), main effect for viral treatment $p = 0.065$, genotype x viral treatment $p = 0.026$. The results of Holm-Sidak post hoc analysis are depicted on the graph.

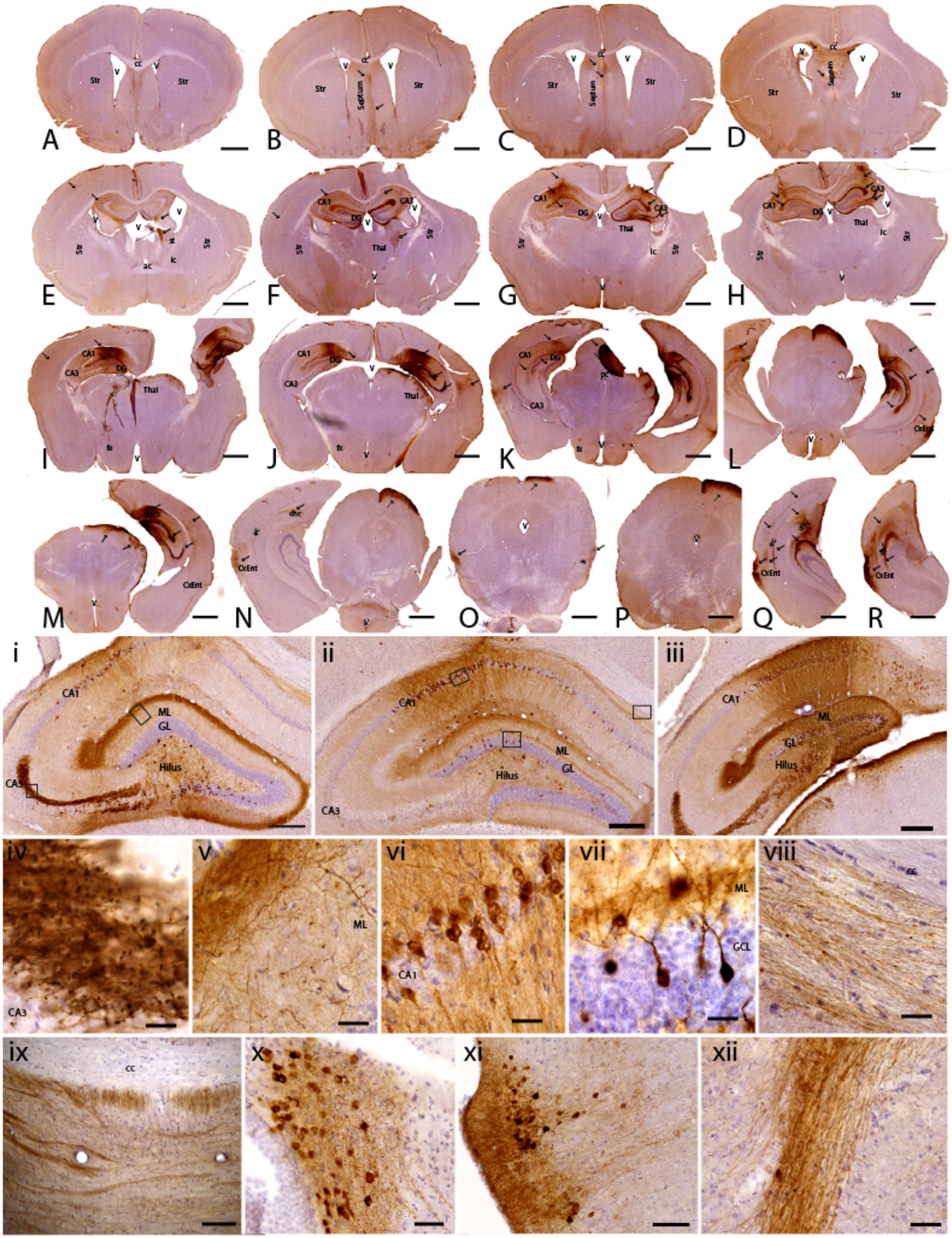
A
B
C
D
E
F
G
H
I
J
K
L
M
N
O
P
Q
R
Str
Septum
CA1
CA3
DG
Thal
CxEnt
i
ii
iii
iv
v
vi
vii
viii
ix
x
xi
xii
ML
GL
Hilus
GCL
cc

**Fig. S5. Widespread expression of CAV-GFP in a juvenile mouse.**

(**A-R**) GFP immunoreactive cells and fibers in a representative DS mouse injected with CAV-GFP at P21. Brains were processed 1-month post-injection. Background staining is hematoxylin and GFP expression is shown by immunohistochemistry DAB staining (dark brown). The arrows indicate the expression of GFP in fibers and cells from the septum **(B-D**), thalamus **(I-J)**, hippocampal regions **(E-M** and **Q-R)**, and the neocortex **(E-M)**, including entorhinal cortex (**K-N** and **Q-R**).

Scale bars: 1 mm.

**GFP immunoreactivity in different brain regions counterstained with hematoxylin.**

***(i-iii)*** High magnifications of hippocampus from rostro to caudal regions, showing the presence of GFP immunoreactive cells and fibers in CA1, CA3 and DG, along the different layers, including the hilus.

(***iv-vii***) Photomicrographs showing the GFP immunoreactivity in the different areas of the hippocampus.

***(iv)*** Detailed of the boxed area in ***(i)*** at the level of the CA3, showing a high number of fibers.

(***v***) GFP immunoreactive fibers at the level of the stratum molecular, and perforant pathway (boxed region in **i**).

***(vi)*** high magnification of the CA1 (from **ii**) showing GFP immunoreactive cells.

***(vii)*** high magnification of the boxed area in the DG of ***ii***, showing the presence of GFP immunoreactive cells also in the granular cell layer of the DG.

***(viii)*** high magnification of GFP immunoreactive fibers in the corpus callosum (cc).

***(ix)*** GFP immunoreactive fibers in the septum.

***(x)*** GFP immunoreactive cells in the dorsal thalamus.

***(xi)*** GFP immunoreactive cells and fibers in the entorhinal cortex.

***(xii)*** GFP immunoreactive fibers in the in the external capsule.

Scale bars **i-iii**: 250 μm; **iv-viii**: 10 μm; **ix**: 100 μm; **x**: 50 μm; **xi**: 100 μm; **xii**: 50 μm.

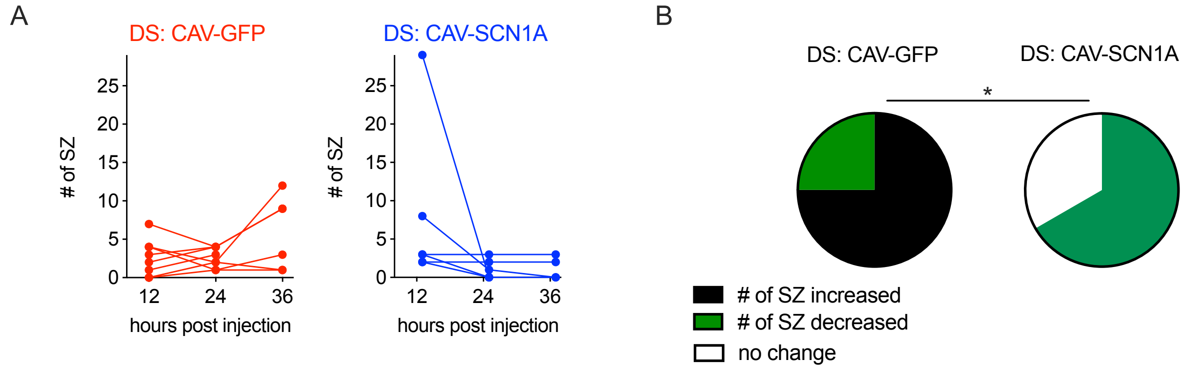


**Fig. S6. Video monitoring of convulsive spontaneous seizures.**

A subset of mice was recorded on video for 12-36 h post-injection.

(**A**) The number of seizures per mouse. The average number of seizures at and 36 h are depicted in Fig. 4B.

(**B**) Change in the frequency of spontaneous seizures over 36 h (Chi-square test $p$ = 0.013).

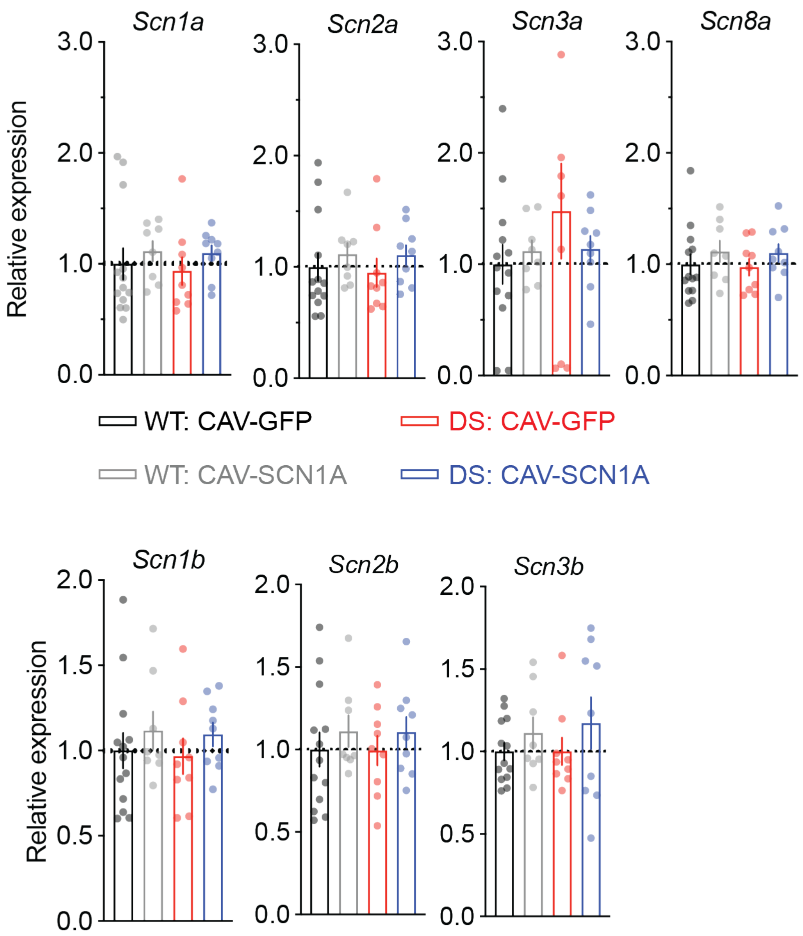


**Fig. S7. Relative expression of voltage gated sodium channels**.

Hippocampi were isolated one month post treatment.

Total RNA was isolated using Purelink RNA mini kit according to the manufacturer's instructions (Thermo Fisher Scientific, Life Technologies, Carlsbald, CA, USA). cDNA was synthesized from 500 ng RNA using Maxima H Minus cDNA synthesis kit (Thermo Fisher Scientific, Life Technologies, Carlsbald, CA, USA). Real-time PCR (qPCR) reactions were performed in triplicates in a final volume of 10 µl with 5 ng of RNA as template using TaqMan gene expression assay (Applied Biosystems, Thermo Fisher Scientific, Life Technologies, Carlsbald, CA, USA ) on StepOnePlus ® real-time PCR system (Applied Biosystems, Thermo Fisher Scientific, Life Technologies, Carlsbald, CA, USA). The following assays were used: *Scn1a (*Mm00450580_m1)*, Scn2a* (Mm01270359_m1)*, Scn3a (*Mm00658167_m1)*, Scn8a (*Mm00488110_m1*), Scn1b* (Mm00441210_m1)*, Scn2b* (Mm01179204_g1)*, Scn3b (*Mm00463369_m1). Two endogenous controls were used: *Gusb (*Mm00446953_m1) and *Tfrc (*Mm00441941_m1). Efficiency of 100%, dynamic range, and lack of genomic DNA amplification were verified for all the assays. WT: CAV-GFP, n = 13; WT: CAV-SCN1A, n = 8; DS: CAV-GFP, n = 9; DS: CAV-SCN1A, n = 9.

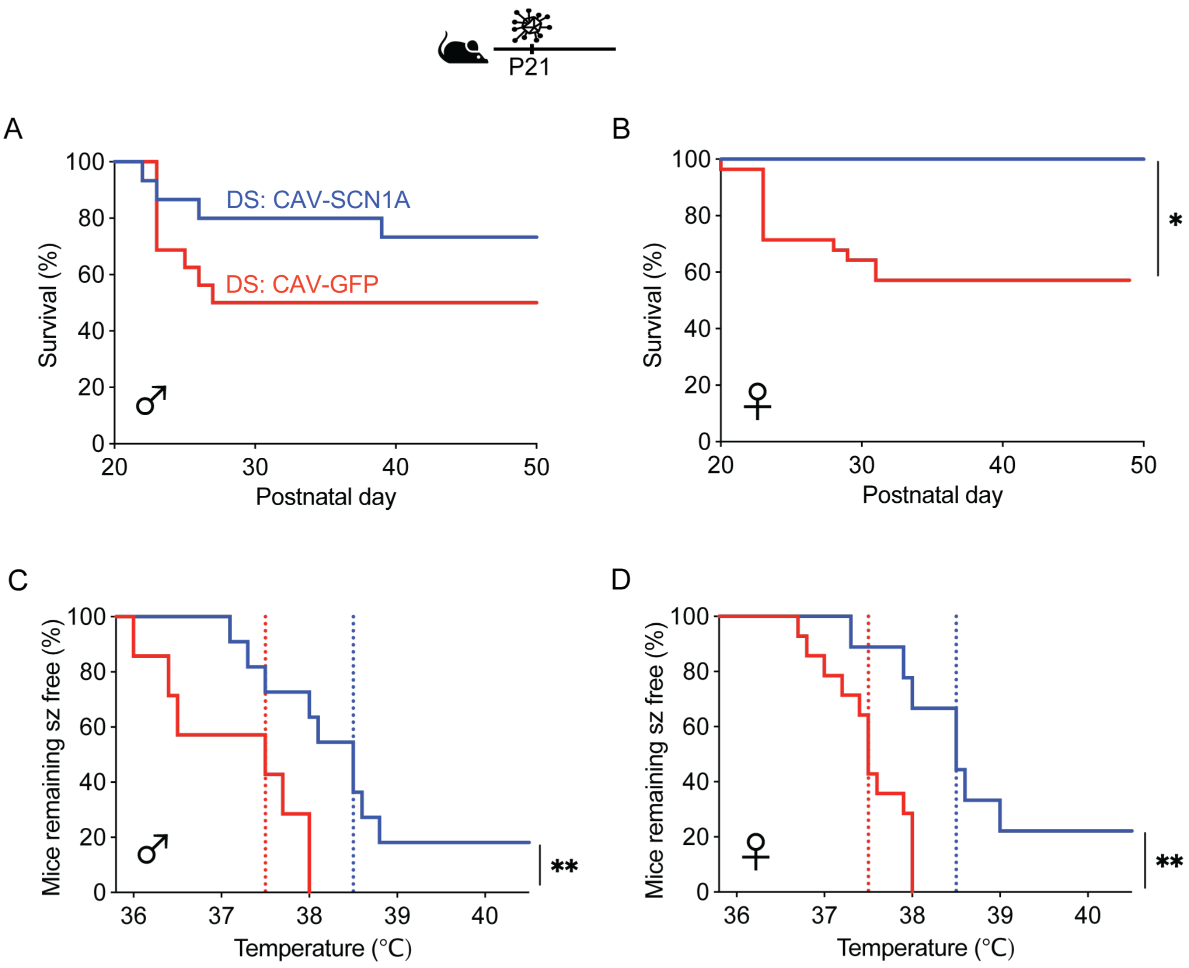


**Fig. S8**. **A similar effect for CAV-SCN1A injection into the thalamus of males and females DS mice.** (**A-B**) Survival curve of males (**A**) and females (**B**) DS mice injected with either CAV-GFP or CAV-SCN1A at P21. DS: CAV-GFP, ♂ n = 16, ♀ n = 28; DS: CAV-SCN1A, ♂ n = 15, ♀ n = 10.

(**C-D**) Males (**C**) and females (**D**) DS mice remaining free of thermally-induced seizures. The dotted lines represent the median seizure temperature. DS: CAV-GFP, ♂ n = 7, ♀ n = 14; DS: CAV-SCN1A, ♂ n = 11, ♀ n = 9. Statistical analysis utilized the Log-rank test. *$p < 0.05$; ** $p < 0.01$.

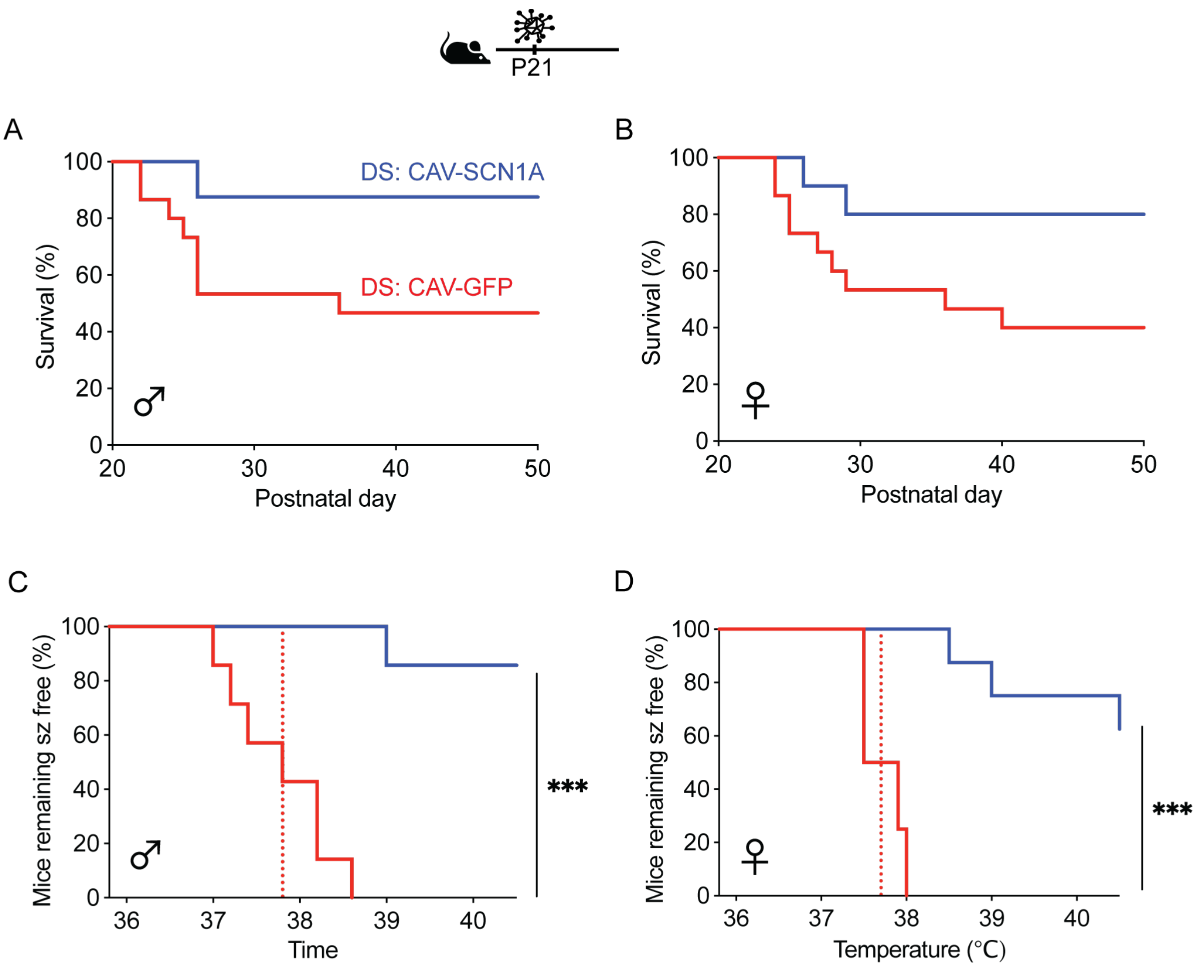


**Fig. S9**. **A similar effect for CAV-SCN1A injection into the thalamus and hippocampus of males and females DS mice.**

(**A-B**) Survival curve of males (**A**) and females (**B**) DS mice injected with either CAV-GFP or CAV-SCN1A at P21. DS: CAV-GFP, ♂ n = 15, ♀ n = 15; DS: CAV-SCN1A, ♂ n = 8, ♀ n = 10.

(**C-D**) Males (**C**) and females (**D**) DS mice remaining free of thermally-induced seizures. The dotted lines represent the median seizure temperature. DS: CAV-GFP, ♂ n = 7, ♀ n = 4; DS:CAV-SCN1A, ♂ n = 7, ♀ n = 8. *** $p < 0.01$.

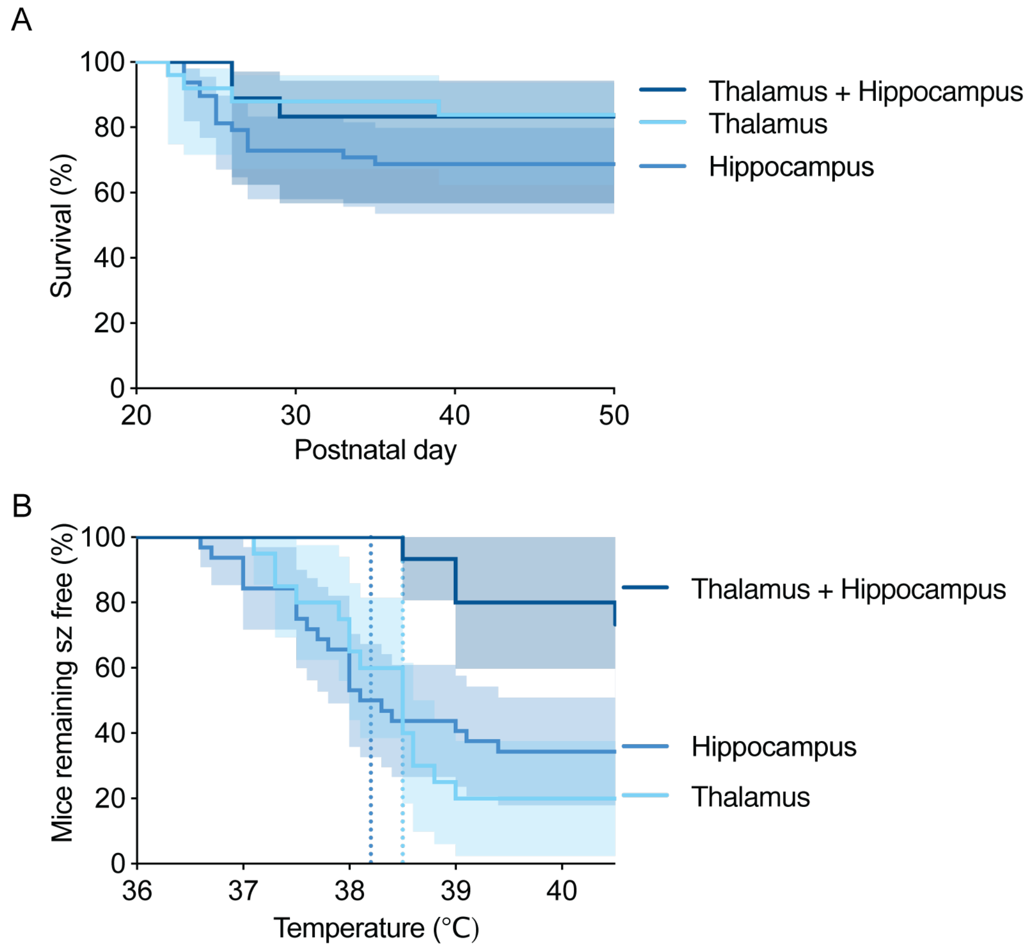


**Fig. S10**. **The effect of CAV-SCN1A injections into the hippocampus, thalamus, or both, in juvenile DS mice.**

(**A**) No statistical difference in the survival of DS mice following CAV-SCN1A injections into the hippocampus, the thalamus, or dual injections to both locations. The solid lines are the same data presented in **Figs. 4A, 6A, 7A**, respectively, and the shaded areas depict 95% confidence intervals.

(**B**) Injection of CAV-SCN1A into the thalamus and hippocampus provides greater protection from thermally-induced seizures. The solid lines are the same data presented in **Figs. 4C, 6B, 7B**, respectively, the shaded areas depict 95% confidence intervals. Note the lack of overlap of 95% confidence intervals following combined injection into the thalamus and the hippocampus, demonstrating greater protection from thermally-induced seizures.
